# Characterization of *C. difficile* strains isolated from companion animals and the associated changes in the host fecal microbiota

**DOI:** 10.1101/822577

**Authors:** R. Thanissery, M.R. McLaren, A. Rivera, Amber D. Reed, N.S. Betrapally, T. Burdette, J.A. Winston, M. Jacob, B.J. Callahan, C.M. Theriot

## Abstract

**Background:** *Clostridioides difficile* is an enteric pathogen historically known to cause hospital associated (HA)-infections in humans. A major risk factor for CDI in humans is antibiotic usage as it alters the gut microbiota and there is a loss of colonization resistance against *C. difficile*. In recent years there has been an increase in community associated (CA)-*C. difficile* infection that does not have the same risk factors as HA-CDI. Potential sources of CA-CDI have been proposed and include animals, food, water, and the environment, however these sources remain poorly investigated. Here, we define the prevalence of *C. difficile* strains found in different companion animals (canines, felines, and equines) to investigate a potential zoonotic link. *C. difficile* strains were identified by toxin gene profiling, fluorescent PCR ribotyping, and antimicrobial susceptibility testing. 16s rRNA gene sequencing was done on animal feces to investigate the relationship between the presence of *C. difficile* and the gut microbiota in different hosts.

**Results:** Here, we show that *C. difficile* was recovered from 20.9% of samples (42/201), which included 33 canines, 2 felines, and 7 equines. Over 69% (29/42) of the isolates were toxigenic and belonged to 14 different ribotypes, with overlap between HA- and CA-CDI cases in humans. The presence of *C. difficile* results in a shift in the fecal microbial community structure in both canines and equines. Commensal Clostridia *C. hiranonis* was negatively associated with *C. difficile* in canines. Further experimentation showed a clear antagonistic relationship between the two strains *in vitro*, suggesting that commensal *Clostridia* might play a role in colonization resistance against *C. difficile* in different hosts.

**Conclusions:** In this study we investigated a potentially important source of *C. difficile* transmission: the companion animal population. *C. difficile* carriage was common in dogs, cats, and horses. *C. difficile* isolates from companion animals included many of the same ribotypes known to cause HA- and CA-CDI in humans, and had similar antimicrobial resistance profiles as those isolated from human populations. These data contribute to our understanding of non-hospital exposure to *C. difficile* in the human population and suggest new avenues for reducing *C. difficile* prevalence in companion animals and, perhaps, thereby reducing CA-CDI in humans.

## Introduction

*Clostridioides difficile* is the most common cause of hospital-acquired and antibiotic-associated diarrhea in the United States, resulting in an estimated 29,000 deaths and over $4.8 billion dollars in medical expenses each year [1, 2]. *C. difficile* was first identified as a primary infectious cause of antibiotic-associated pseudomembraneous colitis and fatal colonic disease in humans in the late 1970s [3, 4]. The widespread use of broad-spectrum antimicrobials contributed to *C. difficile* infection (CDI) becoming a significant hospital-acquired (HA) disease in the subsequent decades [3, 5–7]. It is now understood that many cases of CDI are community-associated (CA-CDI) [8–10]. The epidemiological definition of CA-CDI is broad, meaning no documented overnight stay in a healthcare facility in the prior 12 weeks [2]. In a population-based study from Minnesota, CA-CDI accounted for 41% of all CDI cases [11]. Another study from North Carolina reported that CA-CDI occurred in 21 and 46 per 100,000 person-years in Veterans Affairs (VA) outpatients and Durham County populations, respectively [12]. Traditional risk factors seem to be less important for the new population of CA-CDI that is emerging in younger patients with lower rates of antibiotic exposure [11–14]. The prevalence of CA-CDI cases threatens to undermine the progress being made in controlling CDI.

The sources from which humans acquire CA-CDI are not yet well understood, but companion animals are a strong candidate. Companion animals carry toxigenic and non-toxigenic *C. difficile*, and recent studies show considerable genetic overlap between animal and human *C. difficile* strains [13, 15–21]. *C. difficile* prevalence ranges from 2-40% in the feces of clinically normal and diarrheic dogs and cats [15, 22–26], although its clinical relevance to the animals remains unclear. The close contact between pets and their owners in the 38.4% of US households with dogs and 25.4% with cats [27] makes cross-species transmission far more likely. Only 1.5% of Americans households own a horse, but *C. difficile* is an established cause of diarrhea and colitis in horses [28] with a prevalence of 0-8% in healthy horses [29–33] and 12-90% in diarrheic horses [31, 34, 35] suggesting that *C. difficile* in horses may be more likely to be pathogenic. Even if we assume the lowest reported prevalence rates in dogs, cats and horses, these numbers still indicate that millions of Americans closely interact with companion animals carrying *C. difficile* each year. Other proposed sources of CA-CDI include farm animals, food, water, and the environment, however these potential sources all remain poorly investigated at this time [15-18, 36-39].

The risk factors for CDI or *C. difficile* carriage in companion animals are not well understood. Risk factors associated with CDI in humans have been extensively investigated and include antimicrobial therapy, hospitalization, increasing age, and immunosuppression. It is known that mice, hamsters, and guinea pigs that are used in animal models rapidly develop CDI after antibiotic treatment [40–42]. Some studies in horses show a strong association between antimicrobial therapy and CDI [43, 44], while others indicate no distinct predictors for the disease [45]. Literature on prerequisites for *C. difficile* colonization and infection in dogs and cats mostly show no association with antibiotic therapy [23, 46, 47] while one study showed that treatment with antibiotics was a risk factor for hospital acquired colonization [48].

There is substantial evidence to support the role of shifts in the gut microbiota, especially shifts caused by antibiotics, in the pathogenesis of CDI in humans and animal models [49–51]. The exact molecular mechanisms by which the gut microbiota and their metabolites confer protection are currently being investigated. Several mechanisms such as bile acid transformations [50, 52], competitive exclusion [53] antimicrobial peptides [54], and activation of host immune signaling [55] have been proposed. Therefore, there is growing interest in understanding the interactions between *C. difficile* and the intestinal microbiome. However, taxonomical shifts and microbial interactions associated with *C. difficile* colonization in companion animals have not been explored. Since companion animals show significant genetic overlap of *C. difficile* strains to that of humans, characterization of the *C. difficile* strains circulating in companion animals and identifying their risk factors will help us better understand the epidemiology of this pathogen. Furthermore, defining the fecal microbiota and its interaction with *C. difficile* may identify new taxa associated with protection against *C. difficile* colonization in different hosts. By defining the burden and strains of *C. difficile* prevalent in companion animals we will investigate an understudied source that may contribute to CA-CDI in humans and gastrointestinal disease in companion animals.

In this study, we collected and analyzed 201 fecal samples, from a tertiary hospital, from three companion animals --- canines (dogs), felines (cats), equines (horses) --- and 5 samples from ovines (sheep) as an example farm animal. We determined the prevalence of *C. difficile* in these populations, and *C. difficile* strains isolated from animal stool were characterized in detail by toxin gene profiling, toxin activity testing, ribotyping, and antibiotic susceptibility profiling. Our study revealed widespread prevalence of toxigenic *C. difficile* in companion animals, with significant overlap of ribotypes known to cause HA- and CA-CDI in humans. Using metadata on age, gender, use of antibiotics, and gastrointestinal (GI) health status we determined risk factors associated with *C. difficile* carriage in companion animals. As expected the presence of *C. difficile* correlated with worse GI health status in equines, but not canines.

Fecal microbiota analysis revealed that the commensal *Clostridia Clostridium hiranonis* was negatively correlated to *C. difficile* in canines before and after controlling for antibiotic usage, GI health status, gender, and age. Competition studies with *C. hiranonis* and *C. difficile* show a clear antagonistic relationship between the two strains *in vitro*. We developed further epidemiological and experimental evidence of an interaction between *C. hiranonis* and *C. difficile* in the canine gut, suggesting that commensal *Clostridia* may play a role in colonization resistance against *C. difficile* in different hosts. Our study provides the broadest and most detailed description to date of *C. difficile* across the companion animal species most contacted by Americans. Our results contribute to our understanding of non-hospital exposure to *C. difficile* in the human population and suggest new avenues for reducing *C. difficile* prevalence in companion animals and, perhaps, thereby reducing CA-CDI in humans.

## Materials and Methods

### Collection of animal fecal samples and clinical data

Animal fecal samples submitted for routine microbiological/parasitology diagnostic evaluations to the NC State Veterinary Hospital Microbiology & Molecular Diagnostic Laboratory (MMDL) were randomly selected and collected from December 2016 to October 2017. The included samples were not necessarily representative of GI disease, nor suspected of CDI. The animal species and the fecal sample analysis workflow are presented in Figure 1. A total of 201 samples from canines (n=107), felines (n=17), equines (n= 72), and ovines (n=5) were analyzed. Fecal samples were kept under refrigeration for less than 24 hr, before they were aliquoted, and stored at −80°C until further processing. Metadata on species, breed, age, sex, current antimicrobial therapy (if available), and visit purpose were collected from patient records and discharge comments.

**Figure 1:**
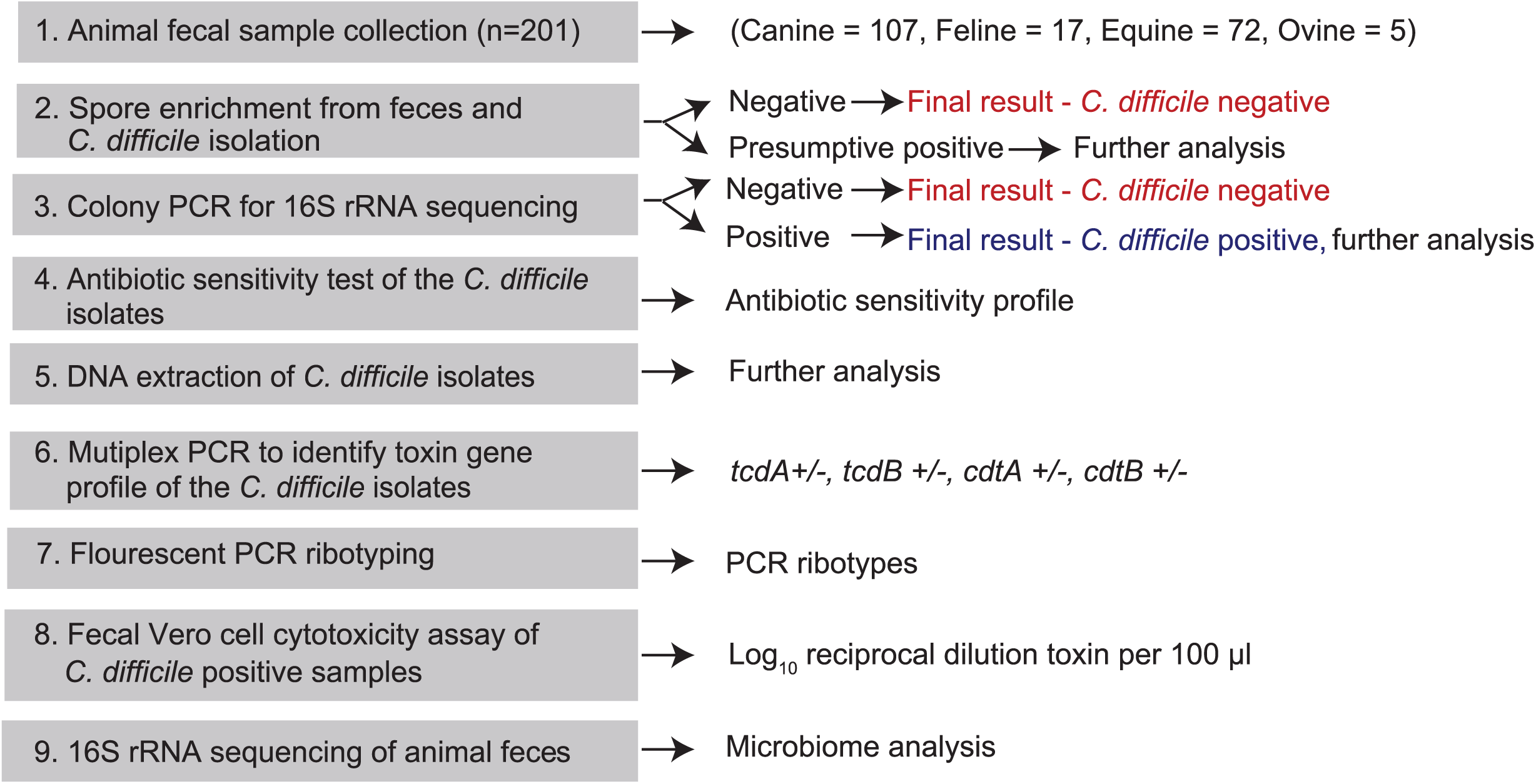
Animal fecal samples analysis workflow. Schematic presenting the workflow of fecal sample collection, overall *C. difficile* prevalence determination, *C. difficile* isolate characterization, and fecal microbiome analysis.

### Spore enrichment from feces and *C. difficile* isolation

*C. difficile* spores were recovered from animal feces as previously described [56]. Fecal samples were thawed and resuspended in 500 µL of pre-reduced phosphate buffer, of which 100 µL were transferred into 5 mL selective taurocholate (Sigma-Aldrich, MO, USA) cycloserine (Fisher Scientific, PA, USA) cefoxitin (Sigma-Aldrich, MO, USA) fructose (Fisher Scientific, PA, USA) broth (TCCFB) for spore enrichment, followed by incubation at 37°C for 24-48 hr anaerobically. The inoculated broth was then inspected for growth by visual turbidity. If turbid, the samples were streaked on a taurocholate cycloserine cefoxitin fructose agar (TCCFA) plate and incubated at 37°C for 24-48 hr anaerobically. The TCCFA plates were inspected for *C. difficile* colony morphology (large flat colonies with appearance of ground glass). The recovered presumptive colonies were further confirmed by direct colony PCR and 16S rRNA gene sequencing (See protocol below). Glycerol stocks (15%) of the confirmed isolates were made and stored at −80°C for downstream analysis. If the colonies present on the TCCFA plates were confluent, and not isolated the colonies were re-streaked onto fresh TCCFA plates and incubated at 37°C for 24-48 hr anaerobically to get isolated colonies. Two to three passages were often needed to obtain pure culture.

### Confirmation of *C. difficile* by direct colony PCR and 16S rRNA gene sequencing

The protocol was adapted from a procedure commonly used for the family Lachnospiraceae [57]. PCR was performed directly on colonies isolated from the TCCFA plates without DNA extraction. The colonies were added to a PCR reaction mixture containing 25 µL of 2X Taq polymerase master mix, universal 16S rRNA primers (Table 1, 1 µL of 5 µM forward primer, 1 µL of 5 µM reverse primer), and Milli-Q water up to 22 µL. The PCR program consisted of 95°C for 5 min; 24 cycles of 95°C for 30 sec, 55°C for 30 sec, and 72°C for 1 min; and final extension at 72°C for 7 min. Positive and negative water controls were used in all runs. The amplified PCR products were analyzed by electrophoresis in a 1.6% agarose gel stained with GelRed. The gels were visualized under UV light, using a Molecular Imager GelDoc ^TM^ XR+ (Bio-Rad). The amplicons were analyzed for nucleotide detection by sanger sequencing. The 16S rRNA gene sequences obtained were processed in BLASTn [58], and analyzed for 100% sequence identity with *C. difficile*.

**Table 1.** Toxin gene profiles of *C. difficile* isolated from different animals

| Toxin gene profile | Animal groups |  |  |  |
| --- | --- | --- | --- | --- |
|  | Canine | Feline | Equine | Total (%) |
| *tcdA, tcdB, cdtA, cdtB* | 0 | 0 | 1 | 1 (2.4) |
| *tcdA, tcdB* | 17 | 1 | 4 | 22 (52.4) |
| *tcdA* | 4 | 0 | 0 | 4 (9.5) |
| *tcdB* | 2 | 0 | 0 | 2 (4.7) |
| None | 10 | 1 | 2 | 13 (30.9) |
| Total | 33 | 2 | 7 | 42 |

### Antimicrobial susceptibility testing

*C. difficile* frozen stocks were cultured on TBHI (Brain heart infusion, Fisher Scientific, PA, USA) supplemented with 100 mg/L L-cysteine (Fisher Scientific, PA, USA) and 0.1% taurocholate (TCA) plates at 37°C for 24 hr. Susceptibility of the *C. difficile* isolates to antibiotics were analyzed using an Etest (bioMerieux, Marcy-l’Étoile, France) as per manufacturer instructions. Briefly an inoculum of 10^5^ cfu/ml of each isolate was applied on a prereduced Brucella agar supplemented with Vitamin K1 and hemin (Hardy Diagnostics, Santa Maria, CA) using a sterile cotton swab. Strips (bioMerieux, Marcy-l’Étoile, France) of cefotaxime (cephalosporin), clindamycin (lincosamides), ciprofloxacin (Fluoroquinolones), levofloxacin (Fluoroquinolones), vancomycin and metronidazole were applied to each plate and incubated at 37°C for 24 hr anaerobically. The test results were interpreted using human CLSI (The Clinical and Laboratory Standards Institute) breakpoints shown in table 5 [59].

### Toxin profiling using Multiplex PCR to identify toxin genes

A multiplex PCR assay described in Presson et al. [60] was used for the detection of *C. difficile* toxin genes A (*tcdA*), and B (*tcdB)*, as well as binary toxins (*cdtA* and *cdtB*), with 16S rRNA genes used as an internal PCR control. Additionally, the following controls were used in each run i) *C. difficile* R20291 (a toxigenic strain with a toxin gene profile of *tcdA* positive, *tcdB* positive, *cdtA/B* positive), *C. difficile* F200 a nontoxigenic strain with a toxin gene profile *tcdA* negative, *tcdB* negative, *cdtA/B* negative [61], and water control. The PCR reaction mixture contained 25 µL of 2X Taq polymerase master mix (New England Biolabs, MA, USA; 1X containing 10 mM Tris-HCl, 50 mM KCl, 1.5 mM MgCl2, 0.2 mM dNTPs, 5% Glycerol, 0.08% IGEPAL® CA-630, 0.05% Tween® 20, 25 U/ml Taq DNA Polymerase), 5 µL of genomic DNA, and 12 primers used at the concentration described in Table 1. Genomic DNA was prepared from *C. difficile* grown overnight on BHI plates. A loop full of cells was harvested from the plates, and subsequently DNA was extracted using a DNeasy UltraClean Microbial Kit (Qiagen Valencia, CA) as per the manufacturer’s instructions. Thermocycler conditions used were 10 min at 94°C, followed by 35 cycles of 50 sec at 94°C, 40 sec at 54°C and 50 sec at 72°C, and a final extension of 3 min at 72°C. The reaction products were separated on a 1.5% agarose (Fisher Scientific, PA, USA) gel and detected by GelRed (VWR International PA, USA), staining. Images were captured using Molecular Imager GelDoc ^TM^ XR+ (Bio-Rad).

### Strain typing using fluorescent PCR ribotyping

Ribotyping was done using fluorescent PCR ribotyping of the 16S and the 23S rRNA intergenic spacer sequence as described previously [62]*. C. difficile* colonies for PCR ribotyping were picked from TBHI plates incubated at 37°C for 24 hr anaerobically. Isolated colonies were sub-cultured into 5 ml BHI broth and additionally incubated overnight at 37°C anaerobically. A 1:10 dilution of this culture was heated at 95°C for 15 min and stored at 4°C until later use as template. The PCR reaction consisted of a 25 µL volume that included 12 μL of Promega PCR Master Mix (Fisher Scientific, PA, USA), 0.5 μL forward primer (GTGCGGCTGGATCACCTCCT) and 0.5 μL 6-carboxyfluorescein (FAM)-labeled reverse primer (56 FAM/CCCTGCACCCTTAATAACTTGACC) that were both adjusted to 10 pmol/μL, 10.5 μL of nuclease free water, and 1 μL of DNA template. The PCR cycling conditions were as follows: 95°C for 10 min followed by 35 cycles of (denaturation – 95°C for 0.5 min, annealing – 55°C for 0.5 min, and elongation – 72°C for 1.5 min), and final elongation of 72°C for 10 min. The fluorescent PCR amplicons obtained were diluted 1:1000 with sterile DNase/RNase free water. The templates were then loaded on to a capillary electrophoresis plate containing a 1:240 ratio of ROX 1000 size standard and Hi-Di Formamide. The resulting chromatograms were analyzed using Applied Biosystems Peak Scanner Software (v. 1.0). The distribution of peaks were then analyzed against a database characterized and validated by Martinson et al., [62] (https://thewalklab.com/tools/) that matches with known ribotypes with the same peaks.

### Vero cell cytotoxicity assay of *C. difficile* positive fecal samples

Fecal *C. difficile* toxin activity of the samples that tested positive for *C. difficile* was measured using a Vero cell cytotoxicity assay [63, 64]. Briefly Vero cells were grown and maintained in DMEM media (Gibco Laboratories, MD, USA) with 10% fetal bovine serum (Gibco Laboratories, MD, USA) and 1% Penicillin streptomycin solution (Gibco Laboratories, MD, USA) in a cell culture incubator (37°C and 5% CO2). Trypsin-EDTA (0.25%; Gibco Laboratories, MD, USA) was added to the cells with a contact time of 2-3 min. Cells that came off the flask surface were gently washed with 1X DMEM media and harvested by centrifugation 1,000 RPM for 5 min. Cells were plated at 1 × 10^4^ cells per well in a 96-well flat bottom microtiter plate (Corning, NY, USA), and incubated overnight at 37°C / 5% CO2. Fecal samples were defrosted on ice and a 1:10 dilution was made using 1X PBS. The samples were then centrifuged at 1,750 RPM for 5 min, and the supernatants were collected from each sample and sterilized by passing them through a single 0.22-µm filter. The sterilized samples were then diluted by 10-fold to a maximum of 10^−6^ using 1X PBS. Sample dilutions were incubated 1:1 with PBS or antitoxin (TechLabs, TX, USA) for 40 min at room temperature after which it was added to the Vero cells. Control containing purified *C. difficile* toxins (A and B; List Biological Labs, CA, USA) and antitoxin were included. Plates were viewed under 200X magnification for Vero cell rounding after an overnight incubation at 37°C / 5% CO2. The cytotoxic titer was defined as the reciprocal of the highest dilution that produced rounding in 80% of Vero cells for each sample.

### 16S rRNA-based bacterial community sequencing using Illumina MiSeq platform

Fecal samples (195) were subjected to community 16S rRNA gene sequencing; 6 samples (2 canines, 2 felines, and 2 equines were excluded due to not having enough sample available or poor quality of the sequencing run. Microbial DNA was extracted from the fecal samples using the Mag Attract Power Microbiome kit (Mo Bio Laboratories, Inc.). A dual-indexing sequencing strategy was used to amplify the V4 region of the 16S rRNA gene [65]. Each 20-µl PCR mixture contained 2 µl of 10X Accuprime PCR buffer II (Life Technologies, CA, USA), 0.15 µl of Accuprime high-fidelity polymerase (Life Technologies, CA, USA), 5 µl of a 4.0 µM primer set, 1 µl DNA, and 11.85 µl sterile nuclease free water. The template DNA concentration was 1 to 10 ng/µl for a high bacterial DNA/host DNA ratio. The PCR conditions were as follows: 2 min at 95°C, followed by 30 cycles of 95°C for 20 sec, 55°C for 15 sec, and 72°C for 5 min, followed by 72°C for 10 min. Libraries were normalized using a Life Technologies SequalPrep normalization plate kit as per manufacturer’s instructions for sequential elution. The concentration of the pooled samples was determined using the Kapa Biosystems library quantification kit for Illumina platforms (Kapa Biosystems, MA, USA). Agilent Bioanalyzer high-sensitivity DNA analysis kit (Agilent CA, USA) was used to determine the sizes of the amplicons in the library. The final library consisted of equal molar amounts from each of the plates, normalized to the pooled plate at the lowest concentration. Sequencing was done on the Illumina MiSeq platform, using a MiSeq reagent kit V2 (Ilumina, CA, USA) with 500 cycles according to the manufacturer’s instructions, with modifications [65]. Sequencing libraries were prepared according to Illumina’s protocol for preparing libraries for sequencing on the MiSeq (Ilumina, CA, USA) for 2 or 4 nM libraries. PhiX and genomes were added in 16S amplicon sequencing to add diversity. Sequencing reagents were prepared according to the Schloss SOP (https://www.mothur.org/wiki/MiSeq_SOP#Getting_started), and custom read 1, read 2 and index primers were added to the reagent cartridge. FASTQ files were generated for paired end reads.

### Community-sequencing bioinformatic analysis

Analysis of the V4 region of the 16S rRNA gene was done in the statistical programming environment R [66] using the package DADA2 [67]. Version 1.8 of the DADA2 tutorial workflow (https://benjjneb.github.io/dada2/tutorial.html) was followed to process the MiSeq data. Forward/reverse read pairs were trimmed and filtered, with forward reads truncated at 230 nt and reverse reads at 160 nt, no ambiguous bases were allowed, and each read required to have less than two expected errors based on their quality scores. Error corrected ASVs were independently inferred for the forward and reverse reads of each sample and then read pairs were merged to obtain amplicon sequence variants (ASVs). Chimeric ASVs were identified and removed. For taxonomic assignment ASVs were compared to the Silva v132 database using the implementation of the RDP naive Bayesian classifier available in the DADA2 R package [68, 69]. A BLASTn web search [58] matched ASV4 to the NCBI 16S reference sequence NR 028611.1 for *C. hiranonis* strain TO-931. ASVs with at least 97% identity to this *C. hiranonis* 16S sequence were then identified using the BLASTn command-line tool [58]. One canine sample and one equine sample received fewer than 30 reads after running DADA2 and so were excluded from subsequent analysis of microbiome profiles. The read depth of the remaining samples ranged from 9338 to 60009 reads.

### C. difficile and C. hiranonis co-culture assay

Competition assays were developed to test the interactions between *C. difficile* strain R20291 and *C. hiranonis* TO-931T. Overnight cultures of both strains were grown individually in BHI supplemented with 100 mg/L L-cysteine for *C. difficile*, and additionally supplemented with 2 uM hemin for *C. hiranonis*. The cultures were grown anaerobically at 37°C. After 14 h of growth, both cultures were subcultured to 1:10 and 1:5 into BHI plus L-cysteine and hemin, and allowed to grow for 3-4 h. Once both the cultures doubled they were back diluted in fresh BHI plus L-cysteine and hemin media to obtain a concentration of 1×10^6^ CFU/mL for 1x, 1×10^7^ CFU/mL for 10x, and 1×10^8^ CFU/ mL for 100x. Competition controls included monocultures of 1x *C. difficile*, 1x *C. hiranonis*, 10x *C. hiranonis*, and 100x *C. hiranonis. C. difficile* and *C. hiranonis* were mixed at 1:1, 1:10, and 1:100 ratios respectively. Appropriate dilutions were plated after 24 h of incubation of all treatments anaerobically at 37 °C. Bacterial enumeration was performed and expressed as Log CFU/mL of culture.

### Statistical analysis

Fisher’s exact test for significance, odds ratios, and confidence intervals for the host metadata-*C. difficile* associations reported in Table 3 were evaluated using Prism version 7.0 for Windows (GraphPad Software, La Jolla, CA, United States). Statistical significance was set at a *p*-value of <0.05 for all analyses (^∗^*p* < 0.05, ^∗∗^*p* < 0.01, ^∗∗∗^*p* < 0.001, ^∗∗∗∗^*p* < 0.0001). All statistical analysis of the microbiota profiles was performed in R; code and data are available on GitHub at https://github.com/rthanis/animal-cdiff. The phyloseq and vegan R packages were used to obtain diversity indices and ordination plots [70, 71]. Associations of animal type and *C. difficile* with alpha diversity were measured by a two-sided Wilcoxon rank sum test, and associations with Bray-Curtis beta diversity were done by permanova using the adonis2 function from the vegan package. Differential-abundance analysis was performed using the ALDEx2 R package [72] and visualized with the ggplot2 R package [73]. Logistic regression of *C. difficile* presence in canines against *C. hiranonis* and other host variables was performed using the brms R package [74, 75] interface to the Bayesian statistical inference software Stan [76].

## Results

### *C. difficile* prevalence, ribotype, and toxin gene profiles vary by animal species

Forty-two *C. difficile* strains were isolated from a total of 201 animal fecal samples submitted to the NC State University Veterinary Hospital MMDL. *C. difficile* was recovered from 30.8% (33/107) of canines, 11.8% (2/17) of felines, 9.7% (7/72) of equines, and no ovines (0/5). To determine the genetic diversity of the *C. difficile* strains circulating in animals we characterized the toxin gene profiles and PCR ribotypes of the isolates. *C. difficile* isolates obtained from all animal groups were subjected to multiplex PCR for detection of genes that encode for toxins A (*tcdA*), B (*tcdB)*, and binary toxins (*cdtA* and *cdtB*). In total, 69% (29/42) of the isolates were positive for at least one of the toxin genes tested, and the rest were non-toxigenic. Five different toxin gene profiles resulted that included 2.4% (1/42) *tcdA tcdB cdtA cdtB*, 52.4% (22/42) *tcdA tcdB*, 9.5% (4/42) *tcdA*, 4.7% (2/42) *tcdB* and 30.9% (13/42) non-toxigenic isolates (Table 1). The gene profile *tcdA tcdB* was the most prevalent accounting for 17/33 of canine, 1/2 of feline, and 4/7 of equine *C. difficile* isolates. Isolates from canines had toxin variant genotypes in addition to the non-variant genotypes. Overall, 18 different known ribotypes were observed and three *C. difficile* isolates had patterns that did not match the established database (Table 2). Different ribotypes were shared between different animal groups. The most frequent canine ribotypes were F014-020 (7/33) and F106 (7/33), followed by FP310 (5/33). The two feline *C. difficile* isolates belonged to two different ribotypes, FP310 and FP501. All seven equine *C. difficile* isolates were also from different ribotypes. One equine isolate, belonging to the ribotype 078-126 encoded all four-toxin genes. Ribotypes associated with human CDI were also detected in some canines and equines, F014-020, F106, 078-126, and F002.

**Table 2.** *C. difficile* PCR ribotypes isolated from different animals

| PCR ribotypes | Animal groups |  |  |  |
| --- | --- | --- | --- | --- |
|  | Canine | Feline | Equine | Total |
| F014-020* | 7 | 0 | 1 | 8 |
| F106* | 7 | 0 | 1 | 8 |
| FP310 | 5 | 1 | 0 | 6 |
| FP313 | 3 | 0 | 0 | 3 |
| F010 | 2 | 0 | 1 | 3 |
| FP415 | 1 | 0 | 0 | 1 |
| F087 | 1 | 0 | 0 | 1 |
| FP418 | 1 | 0 | 0 | 1 |
| FP484 | 1 | 0 | 0 | 1 |
| FP499 | 1 | 0 | 0 | 1 |
| FP407 | 0 | 0 | 1 | 1 |
| 078-126* | 0 | 0 | 1 | 1 |
| F002* | 0 | 0 | 1 | 1 |
| FP501 | 0 | 1 | 0 | 1 |
| Unnamed ribotypes | 2 | 0 | 1 | 3 |
| # of different ribotypes | 10 | 2 | 6 | 18 |
| Total | 31 | 2 | 7 | 40 |

\**C. difficile* ribotype reported to be associated with acute *C. difficile* infection and relapses in humans. Ribotypes separated by hyphen represent similar strains and may belong to either ribotype.

**Table 3.**
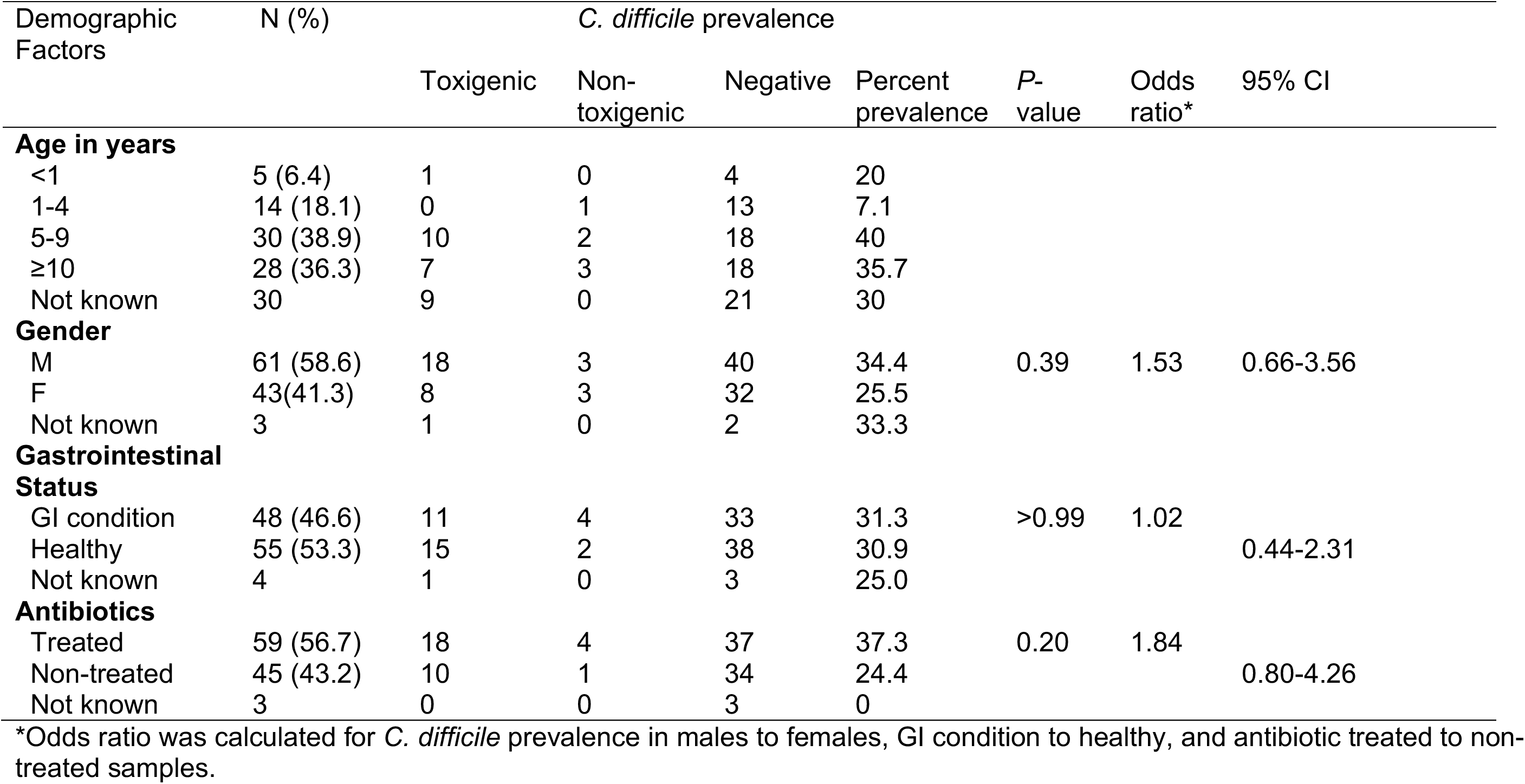
Associations between clinical characteristics and *C. difficile* prevalence outcomes in canines

### Most animals carrying toxigenic *C. difficile* had no detectable toxin activity in their feces

Isolation of toxigenic strains of *C. difficile* from the feces of animals led us to test if there was detectable toxin activity in the fecal samples using a Vero cell cytotoxicity assay. Toxin activity was below the limit of detection in 39 of the 42 fecal samples that were positive for *C. difficile* carriage. In two samples the toxin activity levels were low and only resulted in 50% rounding in the highest concentration (data not shown). These samples contained *C. difficile* isolates belonging to ribotype F106 or F014-020 and belonged to canines, which were treated for non-gastrointestinal related issues. The other sample had a titer of 2 Log10 reciprocal dilution toxin per 100µL/mL fecal sample, and contained *C. difficile* belonging to an unidentified ribotype. This sample also belonged to a canine, which was treated for anorexia and anemia.

### *C. difficile* prevalence in canines was not significantly associated with key demographic factors

The demographic data of canines and equines, which had the most samples, was analyzed further. The key demographic variables including age, gender, gastrointestinal (GI) health status, and antibiotic use in canines and their association with *C. difficile* prevalence are presented in Table 3. The overall proportion of canines with reported GI disease at the time of sampling was 46.3% (48/103). Animals on antibiotics during the time of treatment for which medication data was available was 56.7% (59/104). The study included males (58.6%; 61/104) and females (41.3%; 43/104). A total of 44 different dog breeds with an age range of 3 months to 13 years (median 8 years) were grouped into 4 age categories: <1 yr (6.4%; 5/77), 1-4 yr (18.1%; 14/77), 5-9 yr (38.9%; 30/77), ≥ 10 yr (36.3%; 28/77). A two-sided Fisher’s exact test between *C. difficile* presence and gender, GI health status, or concurrent use of antibiotics in canines all yielded p-values above 0.05 (Table 3). Antibiotic intake appeared moderately positively associated, with an odds ratio of 1.84 (95% confidence interval (0.80, −4.26); p = 0.2). In an exploratory analysis that considered age as a continuous variable, we did observe a higher prevalence of *C. difficile* in dogs between 5 and 11 years and lower in younger and older dogs (Supplementary Figure 1).

### *C. difficile* prevalence in equines is associated with GI health status

The demographic variables and their association with *C. difficile* prevalence in equines are presented in Table 4. In equines, 53.8% (35/65) had symptoms indicative of a GI disorder and 62.1% (36/58) of the reported cases were known to be on antibiotics. The sampled population included males (60.9%; 39/64) and females (39%; 25/64). Equines were grouped into three age categories <2 (9.6%; 5/52), 2-10 (38.5%; 20/52), >10 (51.9; 27/52) and included 25 different breeds. Age was not tested for significance because of the low number of *C. difficile* positives under each category. GI health status was positively associated with *C. difficile* presence (p=0.01 by two-sided Fisher’s exact test; odds ratio 95% confidence interval of (1.66, ∞). All equine samples from which *C. difficile* was recovered belonged to animals that had colic or other GI related issues.

**Table 4.**
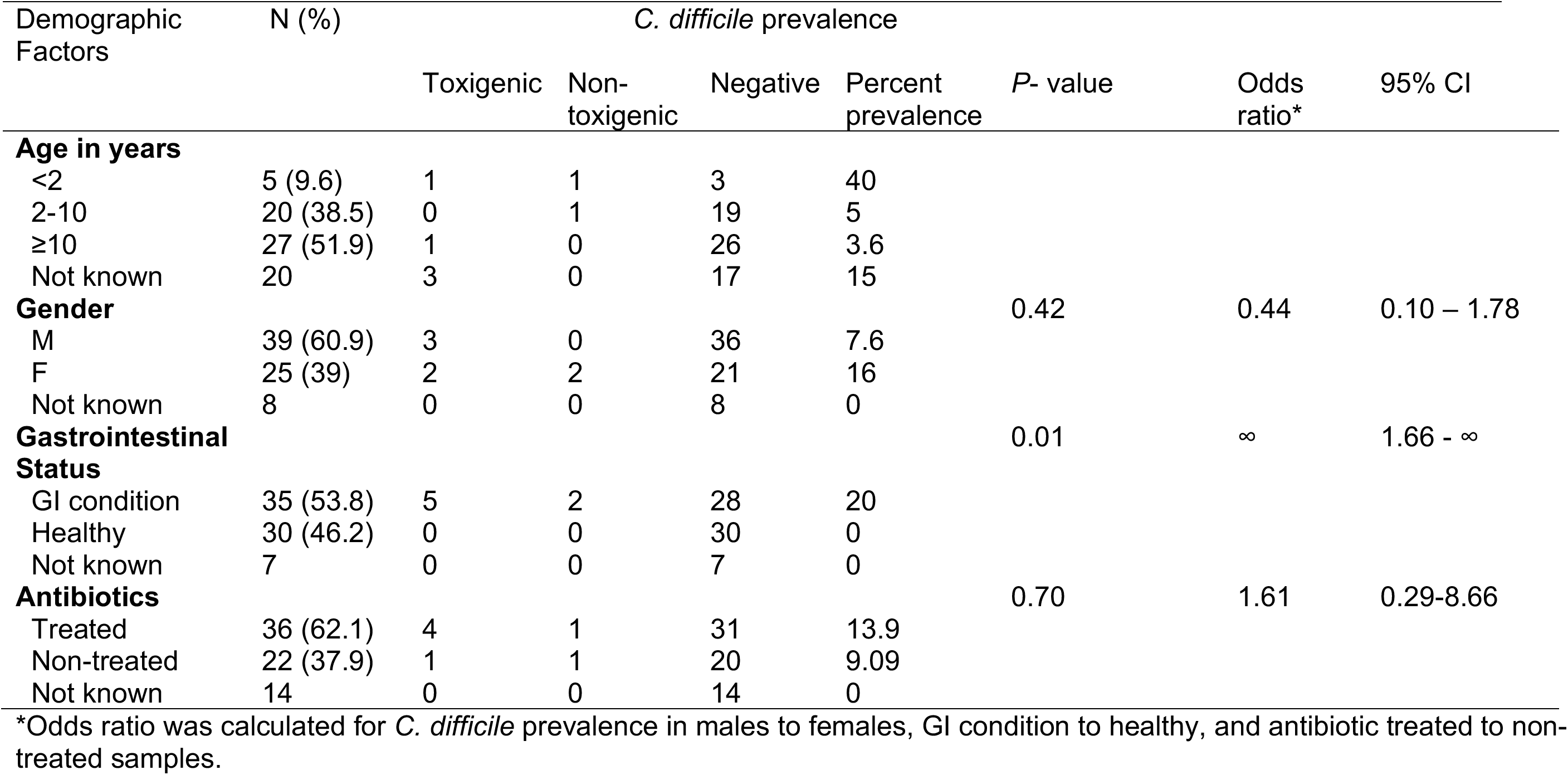
Association between clinical characteristics and *C. difficile* prevalence outcomes in equines

**Table 5.**
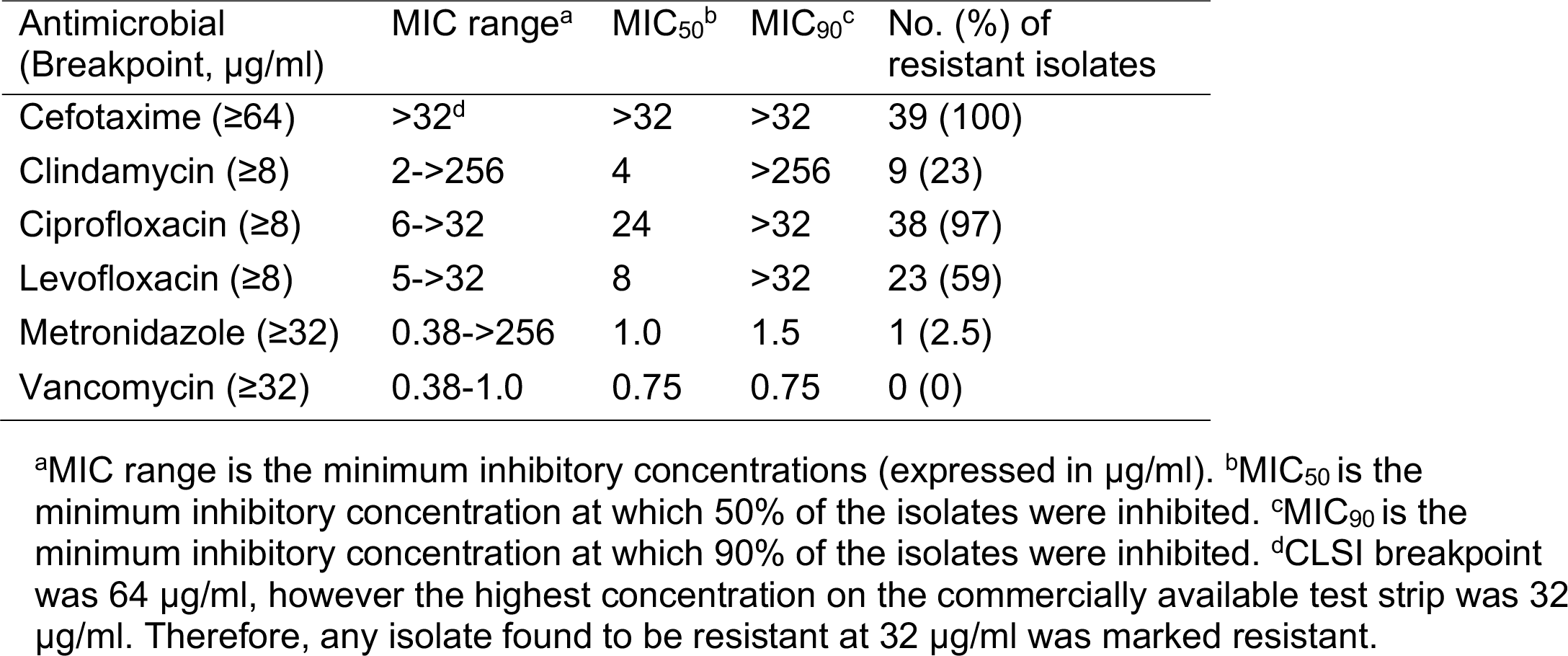
*In vitro* antimicrobial susceptibility of *C. difficile* isolates

### *C. difficile* isolates from companion animals were resistant to antibiotics commonly used in clinical settings, but not front line antibiotics used to treat CDI

To further characterize the *C. difficile* strains isolated from animals, we tested the frequency of resistance to antibiotics used to treat CDI (vancomycin and metronidazole), and those considered as risk factors (cefotaxime, clindamycin, ciprofloxacin, and levofloxacin) for human CDI. All isolates were resistant to cefotaxime and the majority (97%) of them were resistant to ciprofloxacin, whereas resistance to clindamycin and levofloxacin varied (23% and 59% respectively) (Table 5). Only one toxigenic isolate (2.5%, ribotype F014-020) from canine was resistant to metronidazole with a MIC of 512 µg/ml. In all other isolates the MIC for metronidazole remained low (0.38 to 1.5 µg/ml). All *C. difficile* isolates characterized were susceptible to vancomycin.

### *C. difficile* can be detected from animal feces using Illumina 16S rRNA gene sequencing

We sequenced the V4 region of the 16S rRNA gene and processed the resulting sequencing reads with DADA2 [67] to obtain amplicon sequence variants (ASVs). In contrast to traditional 97% sequence similarity OTUs, ASVs have single-nucleotide resolution, which allows species-level classification for some species by exact matching of V4 amplicon sequences [77]. The DADA2 species-classification algorithm identified three *Clostridioides difficile* ASVs (ASV82, ASV1962, and ASV2073) that exactly matched reference *C. difficile* 16S sequences in the Silva database. Two equine samples contained all three ASVs, while all other samples with *C. difficile* contained only ASV82. According to the rrnDB [78] *C. difficile* has 12 copies of the 16S gene and so it is possible that the *C. difficile* strain found in these two equine samples contains all three ASVs at different 16S copies. An additional *Clostridioides* ASV (ASV843) with unknown species identification had 2 nucleotide mismatches from ASV82 and was detected in 3/5 ovine samples and no other animals.

We compared the detection of *C. difficile* via community 16S sequencing to that from the spore-enrichment lab assay. 16S sequencing had good positive predictive value for the results of the lab assay, especially in canines where 19/21 of the samples in which *C. difficile* was detected by 16S sequencing were also positive for *C. difficile* presence in the lab assay. The lab assay detected more *C. difficile* positive samples overall (39 vs. 31) and particularly in canines (31 vs. 21), but each assay detected *C. difficile* in some samples the other assay did not (Table 6). For all subsequent analyses of the effect of *C. difficile* presence on the microbiota, we therefore define samples as *C. difficile* positive if *C. difficile* was detected by the laboratory testing or if ASV82 was detected in the 16S rRNA data.

**Table 6.** Comparison between the detection of *C. difficile* via community 16S sequencing and the spore-enrichment lab assay

|  |  | 16S - | 16S + | Total |
| --- | --- | --- | --- | --- |
| Canine | Lab - | 71 | 2 | 73 |
|  | Lab + | 12 | 19 | 31 |
|  | Total | 83 | 21 | 104 |
| Equine | Lab - | 56 | 6 | 62 |
|  | Lab + | 4 | 3 | 7 |
|  | Total | 60 | 9 | 69 |
| Feline | Lab - | 13 | 1 | 14 |
|  | Lab + | 1 | 0 | 1 |
|  | Total | 14 | 1 | 15 |
| Ovine | Lab - | 5 | 0 | 5 |
|  | Lab + | 0 | 0 | 0 |
|  | Total | 5 | 0 | 5 |
| Total | Lab - | 145 | 9 | 154 |
|  | Lab + | 17 | 22 | 39 |
|  | Total | 162 | 31 | 193 |

The grey cells denote samples for which *C. difficile* was detected by only one assay. Only animals for which 16S data is available are included.

### Differences in the fecal microbiota were associated with animal type and *C. difficile* prevalence

We measured alpha diversity of fecal microbiota samples using inverse Simpson index and evaluated differences between animal type and differences between *C. difficile* positive and negative cohorts within animal types. The distributions of the inverse Simpson index at three microbial taxonomic ranks (Family, Genus, and ASV) are presented in Figure 2A. At each taxonomic rank, canine and feline samples had similar diversities to each other, as did equine and ovine samples, with canine and feline samples having lower diversities than equine and ovine samples. We considered whether *C. difficile* presence was associated with alpha diversity within a host animal type, for the two animals (canines and equines) with larger sample sizes (Supplementary Figure 2). *C. difficile* presence was not associated with genus-level inverse-Simpson diversity in canines (p = 0.84, two-sided Wilcoxon rank sum test; non-parametric 95% confidence interval for the median effect of *C. difficile* presence (−1.1, 1.2) but was associated with lower genus-level inverse Simpson diversity in equines (p = 0.019, two-sided Wilcoxon rank sum test; non-parametric 95% confidence interval for the median effect of *C. difficile* presence (−7.0, −0.75), with similar patterns observed for family- and ASV-level diversity (Supplemental Figure 2).

**Figure 2:**
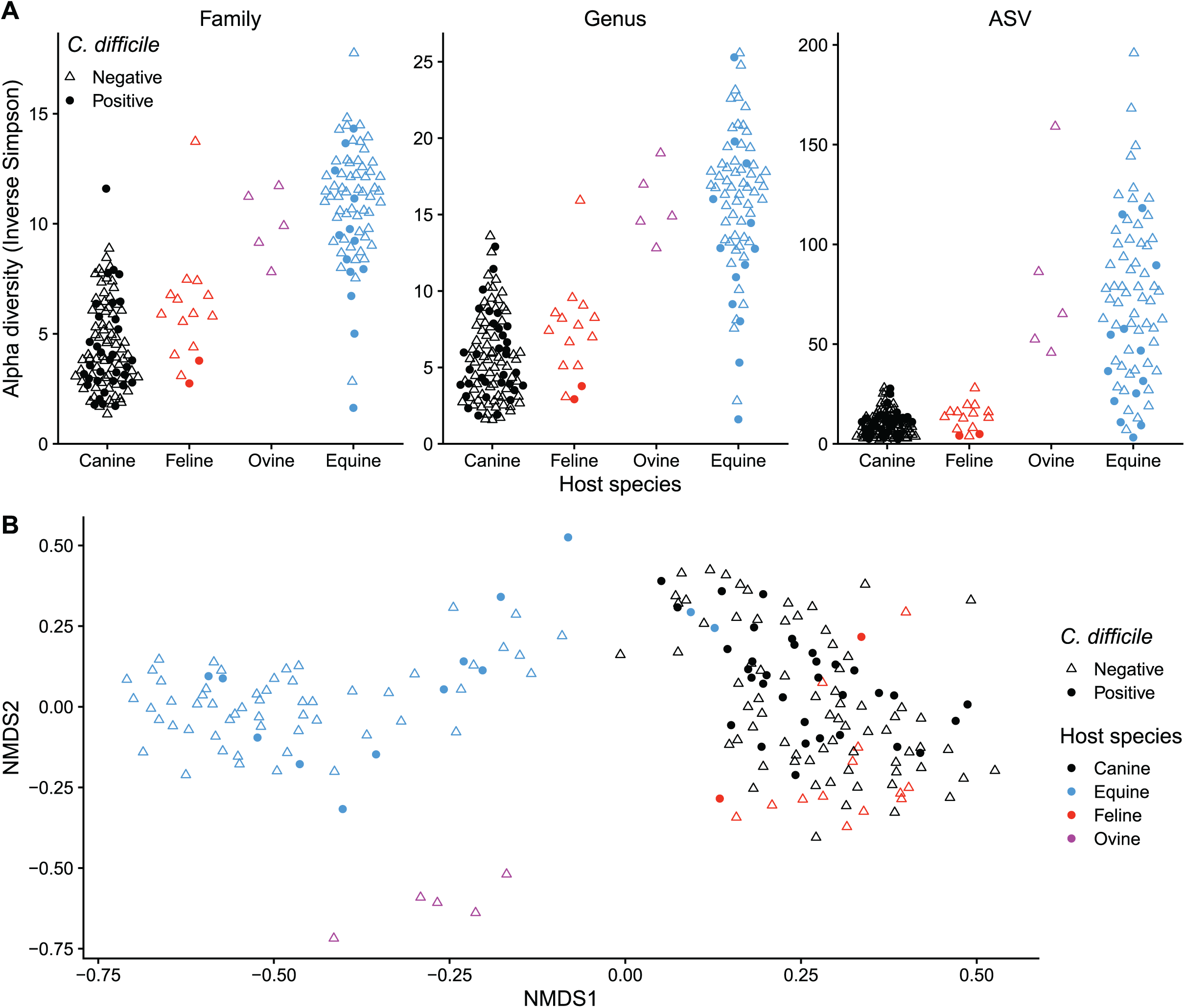
The fecal microbiota associated with *C. difficile* prevalence in animals. A) Alpha diversity in *C. difficile* positive (circles) and negative (open triangle) samples in canines, felines, ovines, and equines. The distribution of inverse Simpson index is presented for different animal groups at the family, genus, and ASV levels. B) Evaluation of beta-diversity in different animal groups that were *C. difficile* positive or negative. Using unsupervised clustering the nonmetric multidimensional scaling (NMDS) illustrates the dissimilarity indices via Bray-Curtis distances between the bacterial communities from animal feces.

To elucidate factors related to the differences and similarities between fecal microbial community structures (β diversity) we performed non-metric multidimensional scaling (NMDS) based on Bray-Curtis dissimilarities between samples at the ASV level. The samples clustered based on animal group (Figure 2B). The first axis separates the canines and felines from the equines and ovines, and the second axis separates the ovines from the equines. No effect of *C. difficile* presence was visually apparent in this cross-species NMDS plot. The presence of *C. difficile* resulted in a statistically significant but weak shift in the fecal microbial community structure in both canines (permonova, *p* = 0.0021, R^2^ = 0.026) and equines (permonova, *p* = 0.00028, R^2^ = 0.030). This result indicates that the degree to which the structure of the whole fecal microbial community could be explained by *C. difficile* presence or absence was low.

### *C. hiranonis* is negatively correlated with *C. difficile* presence in canines

Next, we sought to determine whether the presence of *C. difficile* was associated with specific changes in canine fecal microbial communities. We focused on canines due to the high sample number and the high *C. difficile* prevalence. We performed a compositional principle-components analysis (PCA) [71] to visualize the variation in community composition among canine samples and the ASVs driving this variation. The resulting sample ordination shows a weak but apparent association of *C. difficile* presence with overall community structure (Figure 3A). To determine which ASVs drive this association, we performed a differential-abundance analysis against *C. difficile* presence with the ALDEx2 R package [79]. For each ASV, ALDEx2 reports an effect size estimating the difference in the taxon’s centered-log-ratio (a measure of relative abundance) between groups (*C. difficile* positive vs. negative samples) divided by the difference within groups. Figure 3B shows the ASVs with the largest ALDEx2 effect sizes on the taxa PCA ordination that match the sample ordination of Figure 3A. Visualizing the relative abundances of the top positive and negatively associated ASVs indicates that many of the top associations are largely driven by differential prevalence (presence/absence) (Figure 3C).

**Figure 3:**
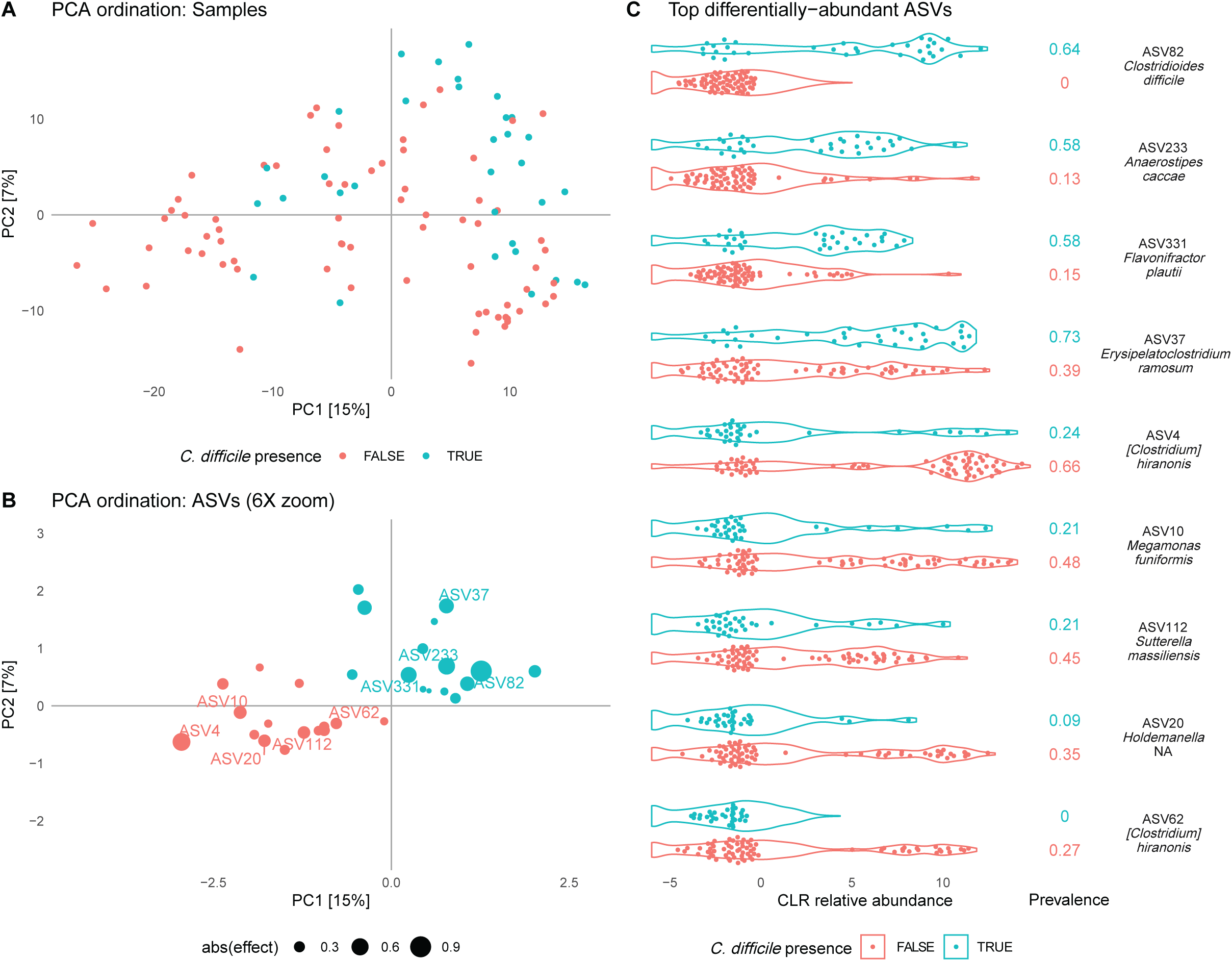
Community composition differences between *C. difficile* positive and negative canine samples using a centered-log-ratio (CLR) transform of the ASV abundances. Community differences were analyzed by principal components analysis (PCA) and differential-abundance analysis (with ALDEx2) of the CLR-transformed abundances. A) and B) show the first two principal components (PCs), which explain 15% and 7% of the variance. Panel A shows the samples and Panel B shows the top ten positive and negatively associated taxa detected by ALDEx2. C) shows the CLR relative abundance for a subset of differentially abundant ASVs in *C. difficile* positive and negative samples. These ASVs are the 4 with the largest positive effect size (ASV82, ASV233, ASV331, and ASV37), the 4 with the largest negative effect size (ASV4, ASV10, ASV20, and ASV112), and the lower-prevalence *C. hiranonis* ASV (ASV62). Violin plots show the estimated distribution of the CLR values in *C. difficile* positive and negative samples, accounting for uncertainty in each sample due to multinomial sampling error during sequencing. Points indicate the mean CLR value for each sample. The numbers to the right of the violin plots indicate the prevalence of the ASV.

The largest negative effect size reported by ALDEx2 is that of ASV4. Although this ASV was not assigned to a species from the Silva database, we used BLAST to determine that it exactly matches the 16S V4 region of [*Clostridium*] *hiranonis* strain TO-931. We therefore sought to further understand the relationship between [*Clostridium*] *hiranonis* with *C. difficile* in our samples. There are 23 ASVs with >99% similarity to *C. hiranonis* strain TO-931; these are ASV4 and 22 additional ASVs that differ from ASV4 by 1 bp. These 22 ASVs have lower prevalence than ASV4 and always appear alongside but have lower abundance than ASV4, suggesting a variety of strains of *C. hiranonis* that carry ASV4 and these other variants at additional copies of the 16S gene. Considering these other ASVs allowed us to identify a possible strain of *C. hiranonis* that may be particularly negatively associated with *C. difficile*. ASV62 appears in 19/52 (27%) of *C. difficile* negative samples but 0/33 of *C. difficile* positive samples (Figure 3C). Bayesian logistic regression of *C. difficile* presence against the presence of ASV4 with or without ASV62 along with sample variables demonstrates that the negative association of *C. difficile* with *C. hiranonis* remains after controlling for antibiotics usage, GI health status, gender, and age and that *C. hiranonis* carrying ASV62 potentially has a much stronger negative effect on *C. difficile* presence than other *C. hiranonis* strains (ASV4 alone: odds-ratio has a 90% credible interval of 0.17–0.57; ASV4 with ASV62: odds-ratio has 90% credible interval of 0.005–0.12). *C. hiranonis* in felines was also negatively associated with *C. difficile* (p = 0.029 by two-sided Fisher’s exact test; 0 of 12 felines with *C. hiranonis* and 2 of 3 felines without *C. hiranonis* are *C. difficile* positive). *C. hiranonis* was only detected in 1 equine sample.

### *C. difficile* suppresses *C. hiranonis* growth in a concentration dependent manner in co-culture *in vitro*

To test the hypothesis that there is an exclusionary relationship between *C. difficile* and *C. hiranonis* in the canine gut, we developed a co-culture *in vitro* assay in rich medium. *C. difficile* and *C. hiranonis* were grown alone (monoculture) and in concert at different concentrations over a 24 hour period in BHI medium (Figure 4). *C. difficile* growth over a 24 hour period was not inhibited by *C. hiranonis* cells at the 1:1 concentration and even when *C. hiranonis* outnumbered *C. difficile* by a factor of 10 (*C. difficile* to *C. hiranonis* ratio), whereas *C. hiranonis* growth was significantly inhibited by 24 hours (Figure 4A and B). It was only when *C. hiranonis* outnumbered *C. difficile* cells by a factor of 100 that *C. hiranonis* growth was not affected at 24 hours (Figure 4C). This provides further evidence that there is a relationship between these two strains, but the mechanism of their interaction still requires further study.

**Figure 4:**
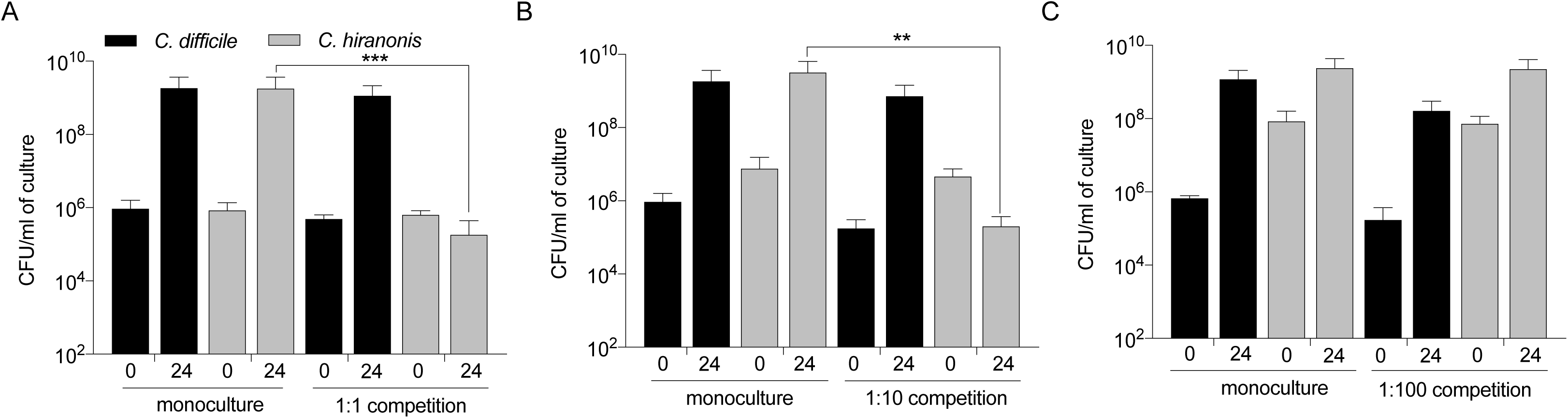
*C. difficile* inhibits *C. hiranonis* growth in co-culture. Co-culture assays in Brain heart infusion media supplemented with 100 mg/L L-cysteine between *C. difficile* and *C. hiranonis*, where cultures were mixed in the ratio of 1:1 (A), 1:10 (B), or 1:100 (C). Colonies were enumerated and expressed as CFU/mL of culture at time points 0 and 24 h for the competition controls (monocultures of *C. difficile* and *C. hiranonis*) and the co-cultures (1:1, 1:10, and 1:100). Data presented represents mean ± SD of at least four experiments in A and B and triplicate experiments in C. Statistical significance between 24 h monoculture and the respective co-culture treatments was determined by Student’s parametric t-test with Welch’s correction (**, p < 0.01; ***, p < 0.001).

## Discussion

The prevalence of *C. difficile* in the dogs, cats, and horses we studied was substantial, and supports the possibility that millions of Americans may be exposed to *C. difficile* each year through contact with companion animals. *C. difficile* was detected in 31% of the canines in our study, which is within the previously reported range (2-40%), although slightly higher than the range (10.5-18.4%) reported from other parts of the U.S, perhaps because our study population was recruited from a tertiary animal hospital [16, 17, 23, 80]. Felines were colonized at 11.8% (2/17) in our study, which is within the previously reported range (2-30%) [16, 22, 24, 26, 48, 81], as was the 9.7% *C. difficile* carriage rate of equines in our study given the varying age and GI health status of the studied population [16, 31, 32]. Significant variation exists in *C. difficile* prevalence rates in companion animals reported by different studies, likely due to various factors such as differences in the type of isolation technique, geographic location, and study population. Nevertheless, the mounting body of evidence is consistent with a large interface between humans and companion animals carrying *C. difficile*.

The *C. difficile* strains isolated from companion animal feces in this study, included ribotypes associated with CDI epidemics in humans. Eighteen different *C. difficile* ribotypes were identified in this study. The most common ribotypes recovered were F014-020 and F106 from both canines and equines. Interestingly, a recent epidemiological survey of human *C. difficile* isolates conducted in the U.S. between 2011 and 2017 reported ribotype 106 as the new dominant ribotype, and the most common isolated from CA-CDI. Ribotype 014-020 was the second most common ribotype among HA-CDI [82]. We also isolated from an equine the 078-126 ribotype that is a known epidemic strain in humans often linked to CA-CDI [82]. The nationwide gradual change in circulating ribotypes in humans and the interspecies sharing of the predominant ribotypes warrant increased utilization of a one health framework that can account for potential zoonotic transmission to understand the changing epidemiology of *C. difficile*.

The clinical relevance of *C. difficile* carriage in dogs and cats remains unclear, but asymptomatic carriage does not preclude transmission to susceptible hosts. In this study, most animals carrying toxigenic *C. difficile* had no detectable toxin activity in their feces via a Vero cell cytotoxicity assay. This could possibly be explained by toxin break down since the fecal samples were not frozen immediately after collection, but is also consistent with largely asymptomatic carriage in dogs and cats. For example, the three dogs that had detectable toxin activity in the feces did not present any atypical fecal consistency or GI-related issues. Asymptomatic toxin carriage has been previously recognized in canines by Weese *et al.* [46], but this does not preclude the possibility of symptomatic CDI developing later or after transmission to a different host species. Asymptomatic carriage where *C. difficile* is able to colonize, proliferate, and produce toxin is also reported in humans, and its role in disease transmission and the mechanism of protection is currently under active investigation [83].

Antimicrobial resistance plays a major role in driving the emergence of epidemic isolates and the associated changes in the epidemiology of *C. difficile* [84]. The low prevalence in our study of resistance to vancomycin and metronidazole, the front-line antibiotics used to treat CDI in humans, is similar to previous reports in humans and animals [32, 82, 85], although there are reports of emerging resistance to metronidazole from other parts of the world [86]. Historically clindamycin, later cephalosporins, and more recently fluoroquinolones are recognized as risk agents for CDI in humans. Resistance to clindamycin was found in 23% of the *C. difficile* isolates in our study. All isolates in our study were resistant to ciprofloxacin a second-generation fluoroquinolone, consistent with patterns in human isolates [87]. Resistance to cefotaxime, a third-generation cephalosporin, was common among all isolates, similar to human studies that report 100% resistance [87, 88]. However, most isolates were sensitive to levofloxacin, a newer third-generation fluoroquinolone. The acquisition of fluoroquinolone resistance by certain epidemic strains (027/BI/NAP1) was the key event linked to their rapid emergence and dissemination in North America [89].

Alterations in the fecal microbiota associated with *C. difficile* carriage in dogs and horses were limited. In canines the alpha diversity was similar between *C. difficile* positive and negative cohorts. However, in equines *C. difficile* prevalence was associated with lower genus-level inverse Simpson diversity very similar to that reported in humans during CDI [51, 90] and asymptomatic carriage [90]. Permanova analysis revealed weak, but significant, differences in community structures between *C. difficile* positive and negative cohorts in canines and equines. Alterations in the fecal microbiota, specifically a decrease in the abundance of specific species and overall diversity, is associated with CDI in humans [51], yet how the gut microbiota of domestic animals changes in the presence of *C. difficile* remains unclear.

The largest negative association between any detected taxa and *C. difficile* was with *C. hiranonis* in dogs and cats. This corroborates the recent report of a similar negative association between *C. difficile* and *C. hiranonis* in opportunistically collected canine fecal samples [91]. *C. hiranonis* is a commensal bacterium that encodes the bile acid inducible (*bai*) operon and is capable of 7α-dehydroxylation of primary bile acids into secondary bile acids. Microbial mediated secondary bile acids are known to play a key role in inhibiting the *C. difficile* life cycle *in vitro* [92, 93]. Secondary bile acid synthesis is regulated by 7α-dehydroxylating gut bacteria from the *Clostridium* cluster XIVa [94], which includes *C. hiranonis* and *C. scindens*. *C. scindens* was associated with partial colonization resistance against *C. difficile* in a mouse model [50], however very little is known about the 7α-dehydroxylating capacity of *C. hiranonis* in the canine gut [95] and needs to be further investigated [96].

The presence of *C. difficile* in the co-culture significantly inhibited *C. hiranonis* growth in a concentration dependent manner *in vitro* using a co-culture system in rich medium. The mechanisms for this suppression are unknown currently, but we hypothesize that competition of nutrients might play a role via the Stickland reaction [95] or *C. difficile* could be making an inhibitory product that inhibits other commensals. Recently, Kang *et al.* found that closely related *C. scindens* produces a tryptophan-derived antibiotic that is able to inhibit *C. difficile* [97]. *C. difficile* was also able to secrete cyclic dipeptides that inhibited *C. scindens*. More work needs to be done using *C. hiranonis* strains isolated from canines, as we were unsuccessful with this isolation, in a medium that mimics the canine gut environment. Nevertheless, the commensal *Clostridium* strains could be a promising probiotic to prevent *C. difficile* in canines and humans.

## Conclusions

In this study we investigated a potentially important source of *C. difficile* transmission: the companion animal population. *C. difficile* carriage was common in dogs, cats, and horses. *C. difficile* isolates from companion animals included many of the same ribotypes known to cause HA- and CA-CDI in humans, and had similar antimicrobial resistance profiles as those isolated from human populations. The large amount of contact between people and companion animals carrying *C. difficile*, as well as the similarities between *C. difficile* isolates from humans and companion animals, are consistent with the possibility of zoonotic transmission. However, it will be critical going forward to develop new studies that can rigorously establish that zoonotic transmission of *C. difficile* is a reality and not just a possibility. Understanding how *C. difficile* is able to colonize different companion animals and what factors are able to prevent colonization will be important for developing novel therapeutics to eradicate *C. difficile* in the animal population, and hopefully thereby decreasing CA-CDI cases in humans.

## Ethics approval and consent to participate

Not applicable.

## Consent for publication

Not applicable.

## Availability of data and material

Raw sequences have been deposited in the Sequence Read Archive (SRA) with SRA accession number: PRJNA562547.

## Competing interests

CMT is a scientific advisor to Locus Biosciences, a company engaged in the development of antimicrobial technologies. CMT is a consultant for Vedanta Biosciences.

## Funding

CMT is funded by the National Institute of General Medical Sciences of the National Institutes of Health under award number R35GM119438 and startup funds from the NCSU CVM.

## Authors’ contributions

RT, AR, ADR, and TB designed and performed the experiments. RT, MRM, NSB, JAW, MJ, BJC and CMT analyzed and interpreted the data. RT, MRM, NSB, and BJC analyzed and displayed the microbiome data. RT, MRM, MJ, BJC and CMT contributed to writing the paper.

All authors edited the manuscript.

## Acknowledgements

None to report.

---

**Supplementary Figure 1: *C. difficile* presence by lab assay versus age in canines.** Each point denotes a sample that was positive via the lab assay or negative. Canine age group in years is listed by circle color.

**Supplementary Figure 2: Alpha diversity versus *C. difficile* presence in canines (A) and equines (B).** Points indicate the inverse Simpson diversity in each sample for each taxonomic rank (Family, Genus, and ASV), with the same color and shape as in Figure 2. Crosses indicate the median and inter-quartile range for that group.

**Supplementary Table 1.**
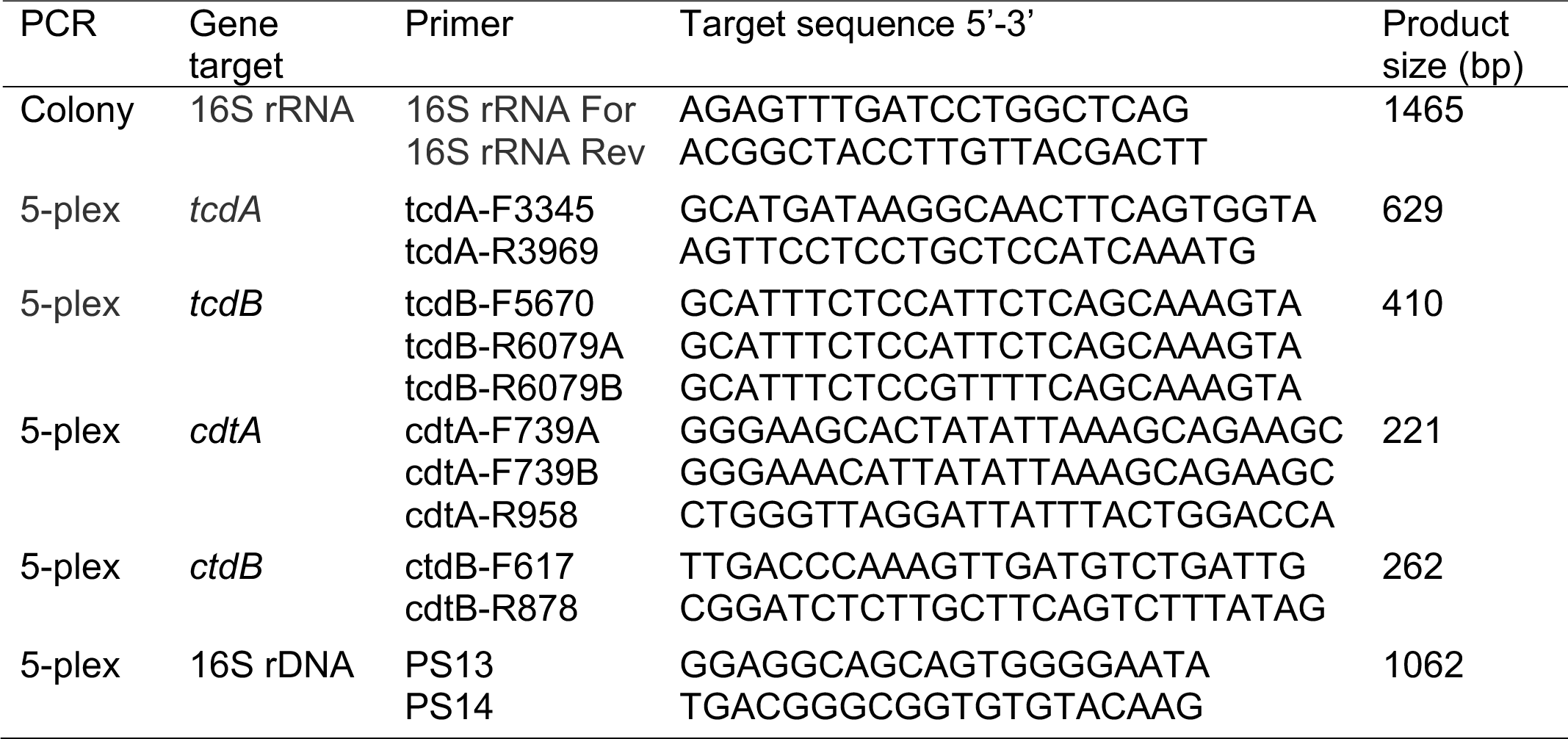
Primer sequences used for colony PCR and 5-plex PCR

## References

1. Lessa FC, Mu Y, Bamberg WM, Beldavs ZG, Dumyati GK, Dunn JR, Farley MM, Holzbauer SM, Meek JI, Phipps EC et al: Burden of *Clostridium difficile* infection in the United States. N Engl J Med 2015, 372(9):825–834.

2. McDonald LC, Gerding DN, Johnson S, Bakken JS, Carroll KC, Coffin SE, Dubberke ER, Garey KW, Gould CV, Kelly C et al: Clinical Practice Guidelines for *Clostridium difficile* Infection in Adults and Children: 2017 Update by the Infectious Diseases Society of America (IDSA) and Society for Healthcare Epidemiology of America (SHEA). Clin Infect Dis 2018, 66(7):e1–e48.

3. Bartlett JG, Chang TW, Gurwith M, Gorbach SL, Onderdonk AB: Antibiotic-associated pseudomembranous colitis due to toxin-producing clostridia. N Engl J Med 1978, 298(10):531–534.

4. Larson HE, Price AB, Honour P, Borriello SP: *Clostridium difficile* and the aetiology of pseudomembranous colitis. Lancet 1978, 1(8073):1063–1066.

5. Pierce Jr P, Wilson Jr R, Silva Jr J, Garagusi V, Rifkin G, Fekety R, Nunez-Montiel O, Dowell Jr V, Hughes J: Antibiotic-associated pseudomembranous colitis: an epidemiologic investigation of a cluster of cases. Journal of Infectious Diseases 1982, 145(2):269–274.

6. Heard S, O’Farrell S, Holland D, Crook S, Barnett M, Tabaqchali S: The epidemiology of *Clostridium difficile* with use of a typing scheme: nosocomial acquisition and cross-infection among immunocompromised patients. The Journal of infectious diseases 1986, 153(1):159–162.

7. Gerding DN, Johnson S, Peterson LR, Mulligan ME, Silva J: *Clostridium difficile*-associated diarrhea and colitis. Infection Control & Hospital Epidemiology 1995, 16(8):459–477.

8. CDC: Severe *Clostridium difficile*-associated disease in populations previously at low risk--four states, 2005. MMWR Morb Mortal Wkly Rep 2005, 54(47):1201–1205.

9. Khanna S, Pardi DS: The growing incidence and severity of *Clostridium difficile* infection in inpatient and outpatient settings. Expert Rev Gastroenterol Hepatol 2010, 4(4):409–416.

10. CDC: Surveillance for community-associated *Clostridium difficile*--Connecticut, 2006. MMWR Morb Mortal Wkly Rep 2008, 57(13):340–343.

11. Khanna S, Pardi DS, Aronson SL, Kammer PP, Orenstein R, St Sauver JL, Harmsen WS, Zinsmeister AR: The epidemiology of community-acquired *Clostridium difficile* infection: a population-based study. Am J Gastroenterol 2012, 107(1):89–95.

12. Kutty PK, Woods CW, Sena AC, Benoit SR, Naggie S, Frederick J, Evans S, Engel J, McDonald LC: Risk factors for and estimated incidence of community-associated *Clostridium difficile* infection, North Carolina, USA. Emerg Infect Dis 2010, 16(2):197–204.

13. Wilcox M, Mooney á, Bendall R, Settle C, Fawley W: A case–control study of community-associated *Clostridium difficile* infection. Journal of Antimicrobial Chemotherapy 2008, 62(2):388–396.

14. Naggie S, Miller BA, Zuzak KB, Pence BW, Mayo AJ, Nicholson BP, Kutty PK, McDonald LC, Woods CW: A case-control study of community-associated *Clostridium difficile* infection: no role for proton pump inhibitors. The American journal of medicine 2011, 124(3):276. e271–276. e277.

15. Weese J, Finley R, Reid-Smith R, Janecko N, Rousseau J: Evaluation of *Clostridium difficile* in dogs and the household environment. Epidemiology & Infection 2010, 138(8):1100–1104.

16. Koene M, Mevius D, Wagenaar J, Harmanus C, Hensgens M, Meetsma A, Putirulan F, Van Bergen M, Kuijper E: *Clostridium difficile* in Dutch animals: their presence, characteristics and similarities with human isolates. Clinical Microbiology and Infection 2012, 18(8):778–784.

17. Stone NE, Sidak-Loftis LC, Sahl JW, Vazquez AJ, Wiggins KB, Gillece JD, Hicks ND, Schupp JM, Busch JD, Keim P: More than 50% of *Clostridium difficile* isolates from pet dogs in Flagstaff, USA, carry toxigenic genotypes. PloS one 2016, 11(10):e0164504.

18. Keel K, Brazier JS, Post KW, Weese S, Songer JG: Prevalence of PCR ribotypes among *Clostridium difficile* isolates from pigs, calves, and other species. Journal of clinical microbiology 2007, 45(6):1963–1964.

19. Arroyo LG, Kruth SA, Willey BM, Staempfli HR, Low DE, Weese JS: PCR ribotyping of *Clostridium difficile* isolates originating from human and animal sources. Journal of medical microbiology 2005, 54(2):163–166.

20. O’Neill G, Adams J, Bowman R, Riley T: A molecular characterization of *Clostridium difficile* isolates from humans, animals and their environments. Epidemiology & Infection 1993, 111(2):257–264.

21. Knight DR, Kullin B, Androga GO, Barbut F, Eckert C, Johnson S, Spigaglia P, Tateda K, Tsai P-J, Riley TV: Evolutionary and genomic insights into *clostridioides difficile* sequence Type 11: A diverse zoonotic and antimicrobial-resistant lineage of global one health importance. mBio 2019, 10(2):e00446–00419.

22. Borriello S, Honour P, Turner T, Barclay F: Household pets as a potential reservoir for *Clostridium difficile* infection. Journal of Clinical Pathology 1983, 36(1):84–87.

23. Struble AL, Tang YJ, Kass PH, Gumerlock PH, Madewell BR, Silva Jr J: Fecal shedding of *Clostridium difficile* in dogs: a period prevalence survey in a veterinary medical teaching hospital. Journal of veterinary diagnostic investigation 1994, 6(3):342–347.

24. Riley T, Adams J, O’neill G, Bowman R: Gastrointestinal carriage of *Clostridium difficile* in cats and dogs attending veterinary clinics. Epidemiology & Infection 1991, 107(3):659–665.

25. Weese JS, Weese HE, Bourdeau TL, Staempfli HR: Suspected *Clostridium difficile*-associated diarrhea in two cats. Journal of the American Veterinary Medical Association 2001, 218(9):1436–1439.

26. Madewell BR, Bea JK, Kraegel SA, Winthrop M, Tang YJ, Silva Jr J: *Clostridium difficile*: a survey of fecal carriage in cats in a veterinary medical teaching hospital. Journal of Veterinary Diagnostic Investigation 1999, 11(1):50–54.

27. Block G, McKenzie B, Yeates J, conditions PiNracoch, $ Pacfevci, AVMA: US pet ownership & demographics sourcebook (2012). J Am Vet Med Assoc 2018, 253(3):264–264.

28. Diab SS, Songer G, Uzal FA: *Clostridium difficile* infection in horses: a review. Veterinary microbiology 2013, 167(1-2):42–49.

29. Frederick J, Giguere S, Sanchez L: Infectious agents detected in the feces of diarrheic foals: a retrospective study of 233 cases (2003–2008). Journal of veterinary internal medicine 2009, 23(6):1254–1260.

30. Medina-Torres CE, Weese JS, Staempfli HR: Prevalence of *Clostridium difficile* in horses. Veterinary microbiology 2011, 152(1-2):212–215.

31. Weese J, Staempfli H, Prescott J: A prospective study of the roles of *Clostridium difficile* and enterotoxigenic *Clostridium perfringens* in equine diarrhoea. Equine veterinary journal 2001, 33(4):403–409.

32. Båverud V, Gustafsson A, Franklin A, Aspan A, Gunnarsson A: *Clostridium difficile:* prevalence in horses and environment, and antimicrobial susceptibility. Equine veterinary journal 2003, 35(5):465–471.

33. McNamara S, Abdujamilova N, Somsel P, Gordoncillo M, DeDecker J, Bartlett P: Carriage of *Clostridium difficile* and Other Enteric Pathogens Among a 4-H Avocational Cohort. Zoonoses and public health 2011, 58(3):192–199.

34. Jones RL, Adney W, Shideler R: Isolation of *Clostridium difficile* and detection of cytotoxin in the feces of diarrheic foals in the absence of antimicrobial treatment. Journal of clinical microbiology 1987, 25(7):1225–1227.

35. Madewell BR, Tang YJ, Jang S, Madigan JE, Hirsh DC, Gumerlock PH, Silva Jr J: Apparent outbreaks of *Clostridium difficile*-associated diarrhea in horses in a veterinary medical teaching hospital. Journal of veterinary diagnostic investigation 1995, 7(3):343–346.

36. Thakur S, Putnam M, Fry PR, Abley M, Gebreyes WA: Prevalence of antimicrobial resistance and association with toxin genes in *Clostridium difficile* in commercial swine. American journal of veterinary research 2010, 71(10):1189–1194.

37. Songer JG, Trinh HT, Killgore GE, Thompson AD, McDonald LC, Limbago BM: *Clostridium difficile* in retail meat products, USA, 2007. Emerging infectious diseases 2009, 15(5):819.

38. Alam MJ, Anu A, Walk ST, Garey KW: Investigation of potentially pathogenic *Clostridium difficile* contamination in household environs. Anaerobe 2014, 27:31–33.

39. Alam MJ, Walk ST, Endres BT, Basseres E, Khaleduzzaman M, Amadio J, Musick WL, Christensen JL, Kuo J, Atmar RL: Community environmental contamination of toxigenic Clostridium difficile. In: Open forum infectious diseases: 2017. Oxford University Press US: ofx018.

40. Lawley TD, Young VB: Murine models to study *Clostridium difficile* infection and transmission. Anaerobe 2013, 24:94–97.

41. Bartlett JG, Onderdonk AB, Cisneros RL, Kasper DL: Clindamycin-associated colitis due to a toxin-producing species of Clostridium in hamsters. Journal of Infectious Diseases 1977, 136(5):701–705.

42. Lowe B, Fox J, Bartlett J: *Clostridium difficile*-associated cecitis in guinea pigs exposed to penicillin. American journal of veterinary research 1980, 41(8):1277–1279.

43. Ruby R, Magdesian KG, Kass PH: Comparison of clinical, microbiologic, and clinicopathologic findings in horses positive and negative for *Clostridium difficile* infection. Journal of the American Veterinary Medical Association 2009, 234(6):777–784.

44. Bäverud V, Gustafsson A, Franklin A, Lindholm A, Gunnarsson A: *Clostridium difficile* associated with acute colitis in mature horses treated with antibiotics. Equine veterinary journal 1997, 29(4):279–284.

45. Weese J, Toxopeus L, Arroyo L: *Clostridium difficile* associated diarrhoea in horses within the community: predictors, clinical presentation and outcome. Equine veterinary journal 2006, 38(2):185–188.

46. Weese JS, Staempfli HR, Prescott JF, Kruth SA, Greenwood SJ, Weese HE: The roles of *Clostridium difficile* and enterotoxigenic *Clostridium perfringens* in diarrhea in dogs. Journal of veterinary internal medicine 2001, 15(4):374–378.

47. Marks SL, Kather EJ, Kass PH, Melli AC: Genotypic and phenotypic characterization of *Clostridium perfringens* and *Clostridium difficile* in diarrheic and healthy dogs. Journal of veterinary internal medicine 2002, 16(5):533–540.

48. Clooten J, Kruth S, Arroyo L, Weese JS: Prevalence and risk factors for *Clostridium difficile* colonization in dogs and cats hospitalized in an intensive care unit. Veterinary microbiology 2008, 129(1-2):209–214.

49. Theriot CM, Koenigsknecht MJ, Carlson Jr PE, Hatton GE, Nelson AM, Li B, Huffnagle GB, Li JZ, Young VB: Antibiotic-induced shifts in the mouse gut microbiome and metabolome increase susceptibility to *Clostridium difficile* infection. Nature communications 2014, 5:3114.

50. Buffie CG, Bucci V, Stein RR, McKenney PT, Ling L, Gobourne A, No D, Liu H, Kinnebrew M, Viale A et al: Precision microbiome reconstitution restores bile acid mediated resistance to *Clostridium difficile*. Nature 2015, 517(7533):205–208.

51. Chang JY, Antonopoulos DA, Kalra A, Tonelli A, Khalife WT, Schmidt TM, Young VB: Decreased diversity of the fecal microbiome in recurrent *Clostridium difficile*—associated diarrhea. The Journal of infectious diseases 2008, 197(3):435–438.

52. Theriot CM, Bowman AA, Young VB: Antibiotic-induced alterations of the gut microbiota alter secondary bile acid production and allow for *Clostridium difficile* spore germination and outgrowth in the large intestine. MSphere 2016, 1(1):e00045–00015.

53. Merrigan MM, Sambol SP, Johnson S, Gerding DN: Prevention of Fatal *Clostridium difficile*–Associated Disease during Continuous Administration of Clindamycin in Hamsters. The Journal of Infectious Diseases 2003, 188(12):1922–1927.

54. Rea MC, Sit CS, Clayton E, O’Connor PM, Whittal RM, Zheng J, Vederas JC, Ross RP, Hill C: Thuricin CD, a posttranslationally modified bacteriocin with a narrow spectrum of activity against *Clostridium difficile*. Proceedings of the National Academy of Sciences 2010, 107(20):9352–9357.

55. Ryan A, Lynch M, Smith SM, Amu S, Nel HJ, McCoy CE, Dowling JK, Draper E, O’Reilly V, McCarthy C: A role for TLR4 in *Clostridium difficile* infection and the recognition of surface layer proteins. PLoS pathogens 2011, 7(6):e1002076.

56. Wilson KH, Kennedy MJ, Fekety FR: Use of sodium taurocholate to enhance spore recovery on a medium selective for *Clostridium difficile*. J Clin Microbiol 1982, 15(3):443–446.

57. Reeves AE, Koenigsknecht MJ, Bergin IL, Young VB: Suppression of *Clostridium difficile* in the gastrointestinal tract of germ-free mice inoculated with a murine Lachnospiraceae isolate. Infection and immunity 2012:IAI. 00647–00612.

58. Altschul SF, Gish W, Miller W, Myers EW, Lipman DJ: Basic local alignment search tool. J Mol Biol 1990, 215(3):403–410.

59. CLSI: Methods for antimicrobial susceptibility testing of anaerobic bacteria; Approved Standard, M11-A8, 8th edn. In.: CLSI Wayne, PA: Clinical and laboratory Standards Institute; 2012.

60. Persson S, Torpdahl M, Olsen KE: New multiplex PCR method for the detection of *Clostridium difficile* toxin A (tcdA) and toxin B (tcdB) and the binary toxin (cdtA/cdtB) genes applied to a Danish strain collection. Clin Microbiol Infect 2008, 14(11):1057–1064.

61. Theriot CM, Koumpouras CC, Carlson PE, Bergin, II, Aronoff DM, Young VB: Cefoperazone-treated mice as an experimental platform to assess differential virulence of *Clostridium difficile* strains. Gut Microbes 2011, 2(6):326–334.

62. Martinson JN, Broadaway S, Lohman E, Johnson C, Alam MJ, Khaleduzzaman M, Garey KW, Schlackman J, Young VB, Santhosh K et al: Evaluation of portability and cost of a fluorescent PCR ribotyping protocol for *Clostridium difficile* epidemiology. J Clin Microbiol 2015, 53(4):1192–1197.

63. Winston JA, Thanissery R, Montgomery SA, Theriot CM: Cefoperazone-treated Mouse Model of Clinically-relevant *Clostridium difficile* Strain R20291. J Vis Exp 2016(118).

64. Thanissery R, Zeng D, Doyle RG, Theriot CM: A Small Molecule-Screening Pipeline to Evaluate the Therapeutic Potential of 2-Aminoimidazole Molecules Against *Clostridium difficile*. Front Microbiol 2018, 9:1206.

65. Kozich JJ, Westcott SL, Baxter NT, Highlander SK, Schloss PD: Development of a dual-index sequencing strategy and curation pipeline for analyzing amplicon sequence data on the MiSeq Illumina sequencing platform. Appl Environ Microbiol 2013, 79(17):5112–5120.

66. Team RC: R: A Language and Environment for Statistical Computing. In. Vienna, Austria; 2019.

67. Callahan BJ, McMurdie PJ, Rosen MJ, Han AW, Johnson AJ, Holmes SP: DADA2: High-resolution sample inference from Illumina amplicon data. Nat Methods 2016, 13(7):581–583.

68. Quast C, Pruesse E, Yilmaz P, Gerken J, Schweer T, Yarza P, Peplies J, Glockner FO: The SILVA ribosomal RNA gene database project: improved data processing and web-based tools. Nucleic Acids Res 2013, 41(Database issue):D590–596.

69. Wang Q, Garrity GM, Tiedje JM, Cole JR: Naive Bayesian classifier for rapid assignment of rRNA sequences into the new bacterial taxonomy. Appl Environ Microbiol 2007, 73(16):5261–5267.

70. McMurdie PJ, Holmes S: phyloseq: An R Package for Reproducible Interactive Analysis and Graphics of Microbiome Census Data. PLoS ONE 2013, 8:e61217.

71. Oksanen J, Blanchet FG, Friendly M, Kindt R, Legendre P, McGlinn D, Minchin PR, O’Hara RB, Simpson GL, Solymos P et al: vegan: Community Ecology Package. In.; 2019.

72. Fernandes AD, Reid JN, Macklaim JM, McMurrough TA, Edgell DR, Gloor GB: Unifying the analysis of high-throughput sequencing datasets: characterizing RNA-seq, 16S rRNA gene sequencing and selective growth experiments by compositional data analysis. Microbiome 2014, 2(1):15.

73. Wickham H: ggplot2: Elegant Graphics for Data Analysis. 2016.

74. Bürkner P-C: brms: An R Package for Bayesian Multilevel Models Using Stan. Journal of Statistical Software 2017, 80:1–28.

75. Bürkner P-C: Advanced Bayesian Multilevel Modeling with the R Package brms. The R Journal 2018, 10:395.

76. Carpenter B, Gelman A, Hoffman MD, Lee D, Goodrich B, Betancourt M, Brubaker M, Guo J, Li P, Riddell A: Stan: A probabilistic programming language. Journal of statistical software 2017, 76(1).

77. Edgar RC: Updating the 97% identity threshold for 16S ribosomal RNA OTUs. Bioinformatics 2018, 34(14):2371–2375.

78. Stoddard SF, Smith BJ, Hein R, Roller BR, Schmidt TM: rrn DB: improved tools for interpreting rRNA gene abundance in bacteria and archaea and a new foundation for future development. Nucleic acids research 2014, 43(D1):D593–D598.

79. Fernandes AD, Reid JN, Macklaim JM, McMurrough TA, Edgell DR, Gloor GB: Unifying the analysis of high-throughput sequencing datasets: characterizing RNA-seq, 16S rRNA gene sequencing and selective growth experiments by compositional data analysis. Microbiome 2014, 2:15.

80. Cave NJ, Marks SL, Kass PH, Melli AC, Brophy MA: Evaluation of a routine diagnostic fecal panel for dogs with diarrhea. JOURNAL-AMERICAN VETERINARY MEDICAL ASSOCIATION 2002, 221(1):52–59.

81. Al Saif N, Brazier J: The distribution of *Clostridium difficile* in the environment of South Wales. Journal of medical microbiology 1996, 45(2):133–137.

82. Tickler IA, Obradovich AE, Goering RV, Fang FC, Tenover FC, Consortium H: Changes in molecular epidemiology and antimicrobial resistance profiles of *Clostridioides (Clostridium) difficile* strains in the United States between 2011-2017. Anaerobe 2019.

83. Gupta A, Khanna S: Community-acquired *Clostridium difficile* infection: an increasing public health threat. Infection and drug resistance 2014, 7:63.

84. Spigaglia P: Recent advances in the understanding of antibiotic resistance in *Clostridium difficile* infection. Therapeutic advances in infectious disease 2016, 3(1):23–42.

85. Usui M, Suzuki K, Oka K, Miyamoto K, Takahashi M, Inamatsu T, Kamiya S, Tamura Y: Distribution and characterization of *Clostridium difficile* isolated from dogs in Japan. Anaerobe 2016, 37:58–61.

86. Baines SD, O’connor R, Freeman J, Fawley WN, Harmanus C, Mastrantonio P, Kuijper EJ, Wilcox MH: Emergence of reduced susceptibility to metronidazole in *Clostridium difficile*. Journal of antimicrobial chemotherapy 2008, 62(5):1046–1052.

87. Bourgault A-M, Lamothe F, Loo VG, Poirier L, group C-Cs: In vitro susceptibility of *Clostridium difficile* clinical isolates from a multi-institutional outbreak in Southern Quebec, Canada. Antimicrobial agents and chemotherapy 2006, 50(10):3473–3475.

88. Simango C, Uladi S: Detection of Clostridium difficile diarrhoea in Harare, Zimbabwe. Transactions of The Royal Society of Tropical Medicine and Hygiene 2014, 108(6):354–357.

89. He M, Miyajima F, Roberts P, Ellison L, Pickard DJ, Martin MJ, Connor TR, Harris SR, Fairley D, Bamford KB: Emergence and global spread of epidemic healthcare-associated *Clostridium difficile*. Nature genetics 2013, 45(1):109.

90. Zhang L, Dong D, Jiang C, Li Z, Wang X, Peng Y: Insight into alteration of gut microbiota in *Clostridium difficile* infection and asymptomatic *C. difficile* colonization. Anaerobe 2015, 34:1–7.

91. Stone NE, Nunnally AE, Jimenez V, Jr., Cope EK, Sahl JW, Sheridan K, Hornstra HM, Vinocur J, Settles EW, Headley KC et al: Domestic canines do not display evidence of gut microbial dysbiosis in the presence of *Clostridioides (Clostridium) difficile*, despite cellular susceptibility to its toxins. Anaerobe 2019.

92. Thanissery R, Winston JA, Theriot CM: Inhibition of spore germination, growth, and toxin activity of clinically relevant *C. difficile* strains by gut microbiota derived secondary bile acids. Anaerobe 2017, 45:86–100.

93. Winston JA, Theriot CM: Impact of microbial derived secondary bile acids on colonization resistance against *Clostridium difficile* in the gastrointestinal tract. Anaerobe 2016, 41:44–50.

94. Berr F, Kullak-Ublick GA, Paumgartner G, Munzing W, Hylemon PB: 7 alpha-dehydroxylating bacteria enhance deoxycholic acid input and cholesterol saturation of bile in patients with gallstones. Gastroenterology 1996, 111(6):1611–1620.

95. Ridlon JM, Devendran S, Alves JM, Doden H, Wolf PG, Pereira GV, Ly L, Volland A, Takei H, Nittono H: The ‘in vivo lifestyle’of bile acid 7α-dehydroxylating bacteria: comparative genomics, metatranscriptomic, and bile acid metabolomics analysis of a defined microbial community in gnotobiotic mice. Gut microbes 2019:1–24.

96. Ridlon JM, Kang D-J, Hylemon PB: Isolation and characterization of a bile acid inducible 7α-dehydroxylating operon in *Clostridium hylemonae* TN271. Anaerobe 2010, 16(2):137–146.

97. Kang JD, Myers CJ, Harris SC, Kakiyama G, Lee IK, Yun BS, Matsuzaki K, Furukawa M, Min HK, Bajaj JS et al: Bile Acid 7alpha-Dehydroxylating Gut Bacteria Secrete Antibiotics that Inhibit *Clostridium difficile*: Role of Secondary Bile Acids. Cell Chem Biol 2019, 26(1):27–34.e24.

