## Supplemental Figure 1 for "Characterization of *C. difficile* strains isolated from companion animals and the associated changes in the host fecal microbiota"

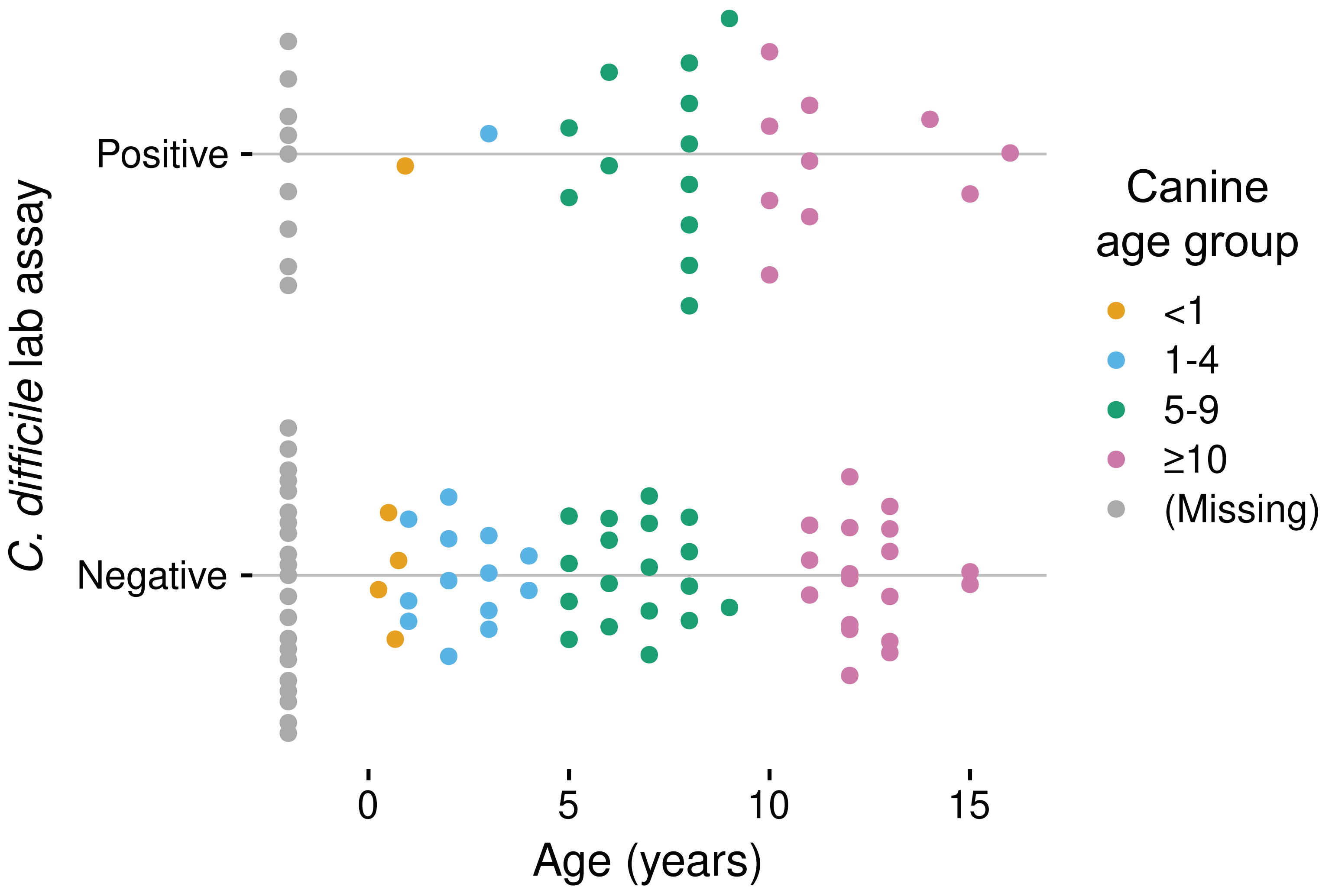


**Supplementary Figure 1: *C. difficile* presence by lab assay versus age in canines.** Each point denotes a sample that was positive via the lab assay or negative. Canine age group in years is listed by circle color.
