## Supplemental Figure 2 for "Characterization of *C. difficile* strains isolated from companion animals and the associated changes in the host fecal microbiota"

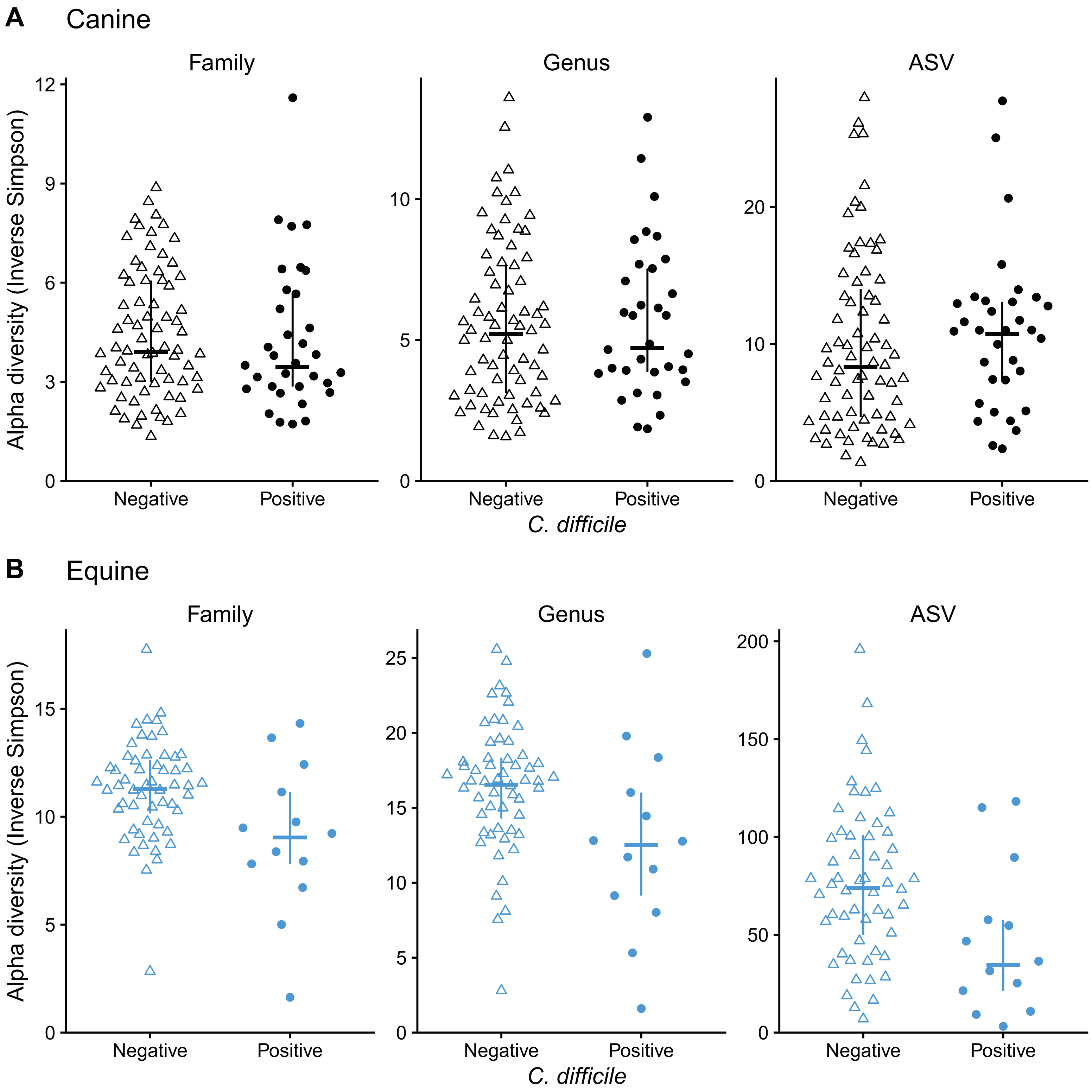


**Supplementary Figure 2: Alpha diversity versus *C. difficile* presence in canines (A) and equines (B).** Points indicate the inverse Simpson diversity in each sample for each taxonomic rank (Family, Genus, and ASV), with the same color and shape as in Figure 2. Crosses indicate the median and inter-quartile range for that group.
